# Hypoalbuminemia Increases Fibrin Clot Density and Impairs Fibrinolysis

**DOI:** 10.1101/2025.02.03.635731

**Authors:** Amanda P. Waller, Katelyn J. Wolfgang, Zachary S. Stevenson, Lori A. Holle, Alisa S. Wolberg, Bryce A. Kerlin

**Author notes:** Corresponding Author: Bryce A. Kerlin, MD Center for Clinical & Translational Research The Abigail Wexner Research Institute at Nationwide Children’s 700 Children’s Dr., W302 Columbus, OH 43205, USA.

## Abstract

**Background:** Hypoalbuminemia is a thrombotic disease risk biomarker. Albumin is a negative acute phase reactant and may thus be an indirect biomarker of thromboinflammation. However, nephrotic syndrome (resulting from non-inflammatory proteinuric glomerular disease) causes both hypoalbuminemia and elevated thrombotic risk. Hypofibrinolysis has been observed in nephrotic syndrome and published data suggest that albumin may directly enhance fibrinolysis. These observations suggest that albumin may directly influence thrombotic risk.

**Objective:** To test the hypothesis that hypoalbuminemia impairs fibrinolysis.

**Methods:** Hypoalbuminemic blood and plasma from nephrotic syndrome rat models and nephrotic patients were analyzed by thromboelastometry, clot lysis assays, fibrin clot turbidity, confocal microscopy, and immunoblotting. Some studies were conducted without vs. with albumin repletion. Plasma from an analbuminemic mutant rat model was used to confirm albumin-dependent observations.

**Results:** Hypoalbuminemia was associated with hypofibrinolysis in nephrotic whole blood. Albumin levels were positively associated with fibrinolysis in both nephrotic rat and patient plasma. Hypoalbuminemia accelerated fibrin clot formation in nephrotic rat plasma. Dense fibrin clots are known to be resistant to fibrinolysis and fibrin clot network density was increased in hypoalbuminemic plasma clots from both nephrotic rats and patients. Clots formed from hypoalbuminemic plasma contained less albumin than controls, and repletion with recombinant albumin to healthy control levels corrected both fibrin network density and fibrinolysis in nephrotic and analbuminemic rat plasma.

**Conclusion:** These data show that albumin directly increases fibrin network porosity and enhances fibrinolysis. Hypoalbuminemia may mechanistically contribute to nephrotic syndrome thrombotic complications and may similarly increase thrombotic risk in other hypoalbuminemic conditions.

## INTRODUCTION

Hypoalbuminemia is a significant predictor of both thrombosis and cardiovascular disease [1-5]. Nephrotic syndrome (NS) is a glomerular disorder characterized by hypoalbuminemia due to massive proteinuria that is associated with a high-risk for life-threatening thrombotic complications [6-9]. Increased thrombotic risk in NS is generally attributed to its well-described acquired hypercoagulability that is caused by deranged protein homeostasis resulting from proteinuria [8]. In addition, however, hypofibrinolysis was described in patients with NS as early as 1974 and may play an underappreciated role in NS-associated thrombotic risk [8, 10-15]. Importantly, decreased clot lysis has been associated with increased thrombotic risk in non-NS cohorts and recent evidence suggests that co-existent hypercoagulability and hypofibrinolysis may synergistically increase thrombosis risk [16, 17]. We previously reported significant hypofibrinolysis in a rat NS model using rotational thromboelastometry and found that the hypofibrinolytic signal was directly correlated with urinary protein and indirectly correlated with plasma albumin [14].

Albumin was previously shown to partially correct hypofibrinolysis in clotted NS patient plasma [11]. The same study demonstrated that fibrin networks from NS patient plasma were more dense, with shorter fibrils and more branch points, a pattern that was partially corrected by albumin supplementation [11]. Other reports, not focused on NS, used plasma, purified fibrinogen, or fibrin monomers to demonstrate that albumin affects fibrin network structure and fibrinolysis [18-23]. However, many of these studies were performed with non-physiologic concentrations of albumin, fibrin(ogen), or both. Many were underpowered, sometimes demonstrating effects in only a single subject. It is thus unclear from these studies if albumin is a biologically important regulator of fibrinolysis.

As a negative acute phase reactant, albumin is a potential thromboinflammation biomarker [3]. However, several studies suggest that albumin may also contribute mechanistically to fibrinolysis [11, 18, 19, 21-23]. Thus, defining the role of albumin in fibrinolytic regulation may increase understanding of thrombotic pathogenesis not only in kidney disease, but also in other diseases associated with hypoalbuminemia [24]. Importantly, hypoalbuminemia is highly prevalent amongst hospitalized patients who are a group that is well-known to be at high risk for thrombosis [24, 25]. We therefore tested the hypothesis that hypoalbuminemia impairs fibrinolysis using both NS and mutant analbuminemic rat models as well as NS patient samples [26].

## MATERIALS AND METHODS

### Hypoalbuminemic Rat Models

All procedures were approved by the Abigail Wexner Research Institute at Nationwide Children’s Hospital Institutional Animal Care and Use Committee (AR13-00027), in accordance with the NIH Guide for the Care and Use of Laboratory Animals. Three hypoalbuminemic rat models were utilized in these studies (**Table S1**). Two of these models are well-established NS models wherein toxin-mediated podocyte injury leads to proteinuric glomerular disease [27, 28]. Hypoalbuminemia develops in both models due to massive albuminuria. We also examined genetically analbuminemic (*Alb*^-/-^) rats that were maintained on an inbred Fischer 344 background [29-32]. These rats have a naturally occurring 7 bp intron deletion that disrupts *Alb* mRNA splicing [29, 30]. As a result, homozygous rats have no detectable plasma or tissue albumin [31, 32]. Blood was collected from wild-type (*Alb*^+/+^), heterozygous (*Alb*^+/-^), and homozygous (Alb^-/-^) littermate rats using the above methods (∼150 g rats, age ∼6 weeks, *n*=5 animals/group). Both male and female rats were studied in the *Alb*^-/-^ model whereas only male rats were considered in the NS models because they had less phenotypic variability in preliminary studies. Administration of puromycin aminonucleoside (PAN), a podocyte-specific toxin, is a well-established dose-dependent rat NS model (PAN-NS) [14, 27, 33, 34]. We previously constructed a repository of plasma samples from Wistar rats with a range of plasma albumin levels by varying the PAN dose [14]. These 5 groups of rats (body weight ∼150 g, age ∼45-50 d) received a single dose of intravenous PAN (0 (sham carrier solution only), 25, 50, 100, or 150 mg/kg; *n*=6 animals/group) on Day 0 [14]. Sham treated rats in this group represent healthy controls and the PAN treated rats were divided into albumin quartiles. We also studied the podocin-promoter driven human Diphtheria Toxin Receptor (*pDTR*) transgene rat NS model (*pDTR*-NS) on an inbred Fischer 344 background [28, 33]. Expression of the *pDTR* transgene in this model is restricted to podocytes. Four groups of homozygous *pDTR* rats (∼150 g, age ∼6 weeks) were treated with intraperitoneal diphtheria toxin (DT) as follows: 0 (sham carrier solution only), 25, 50, or 75 ng/kg on day 0 and compared to a matched group of wild-type Fischer 344 rats treated with 50 ng/kg DT (WT+DT; *n*=5-8 animals/group). This strategy incorporates two types of healthy control: (1) Sham treated *pDTR* rats are presented as the 4^th^ albumin quartile whereas (2) The WT+DT rats are labeled as Controls, since WT rats do not express a DT receptor. Nadir plasma albumin occurs by day 10 in both models [27, 28, 35]. Thus, blood was collected on day 10 from the inferior vena cava into final concentration 0.32% NaCitrate / 1.45 µM Corn Trypsin Inhibitor (CTI; Haematologic Technologies Inc., Vermont, VT, USA), an aliquot of whole blood was used for rotational thromboelastometry (ROTEM) where indicated and the remainder was processed to Platelet Poor Plasma (PPP), within one hour, aliquoted, and frozen at -80°C until analyzed, as described previously [14].

### Columbus Nephrotic Syndrome Cohort

To translate the rat model observations to human hypoalbuminemia, assays were also performed on plasma samples from a previously described cohort of incident pediatric and adult NS patients from whom plasma was collected prior to initiation of NS-specific treatment [36, 37]. This portion of the study was approved by the Nationwide Children’s Hospital Institutional Review Board (IRB12–00290) in reciprocity with The Ohio State University Wexner Medical Center IRB. After written informed consent (and assent when applicable) was obtained from each participant or parent/guardian, blood was collected into final concentration 0.32% sodium citrate / 1.45 μM CTI and processed to PPP as above.

### Plasma Albumin Quantitation

Plasma albumin was quantified in all rat and human samples using the bromocresol purple (BCP) assay (QuantiChrom BCP; BioAssay Systems, Hayward, CA), according to the manufacturer’s instructions, as previously described [34, 36-38].

### Global Hemostasis Assays

Whole blood ROTEM assays were performed within 20 min of specimen collection, using the INTEM (intrinsic pathway activation) assay in mini cups and pins without and with tranexamic acid (TXA; 1 mg per 100 µL of whole blood [39]), on a ROTEM*delta* instrument (Werfen, Barcelona, Spain). ROTEM reports maximal clot firmness (MCF) as the maximal waveform amplitude (in mm). Lysis at 60 minutes (LY60) is the percent decrease in amplitude from MCF at 60 minutes: 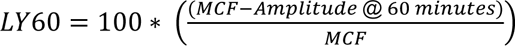 [40, 41]. Thrombin generation assays (TGA) were performed using the Technothrombin TGA kit (Technoclone, Vienna, Austria) on rat PPP diluted 1:1 with TGA buffer and TGA RC Low Reagent, and read on a Spectramax M2 fluorescent plate reader (Molecular Devices, Sunnyvale, California), as previously described [14, 34, 36]. Relative Fluorescence Units (RFU) were converted to thrombin (nM) concentrations using a standard curve and evaluated with Technoclone Evaluation Software.

### Fibrinolysis Assays

Plasma Clot Lysis Assays (CLA) were adapted from a previously reported colorimetric method based upon release of Coomassie brilliant blue R-250 dye from fibrin clots [42]. Briefly, fibrin clot formation was initiated by adding thrombin (20 nM) and calcium chloride (10 mM) to PPP (diluted 1:4 with PBS) in the presence of R-250 dye (1 mg/ml). The plasma was allowed to clot at room temperature for 45 minutes, the resulting clots were washed with 1 mL PBS, and resuspended in 1 mL of a urokinase-type plasminogen activator (uPA) solution (200 IU/mL in PBS). The suspended clot was then gently rotated at room temperature and lysis was determined at 60 minutes by transferring 100 uL of the soluble supernatant fraction into a 96-well microtiter plate wherein absorbance was measured at 540 nm to quantitate R-250 dye released from the clot. Percent total clot lysis was calculated as a percentage of pooled normal plasma (PNP) clot lysis after blank subtraction. In some experiments, the assay conditions were modified to explore potential underlying mechanisms as follows: (a) thrombin (20 nM) without uPA, (b) low thrombin (5 nM) + uPA (200 IU/mL), (c) thrombin (20 nM) + uPA (200 IU/mL) + plasmin (1.2 µM), and (d) thrombin (20 nM) + high uPA (800 IU/mL) with incubation at 37°C on a rotator with endpoint measurement at 24 hrs. In other experiments, CLA was performed with tissue-type plasminogen activator (tPA; 20 nM) or plasmin (1.2 µM) in lieu of uPA or after immunodepleting or supplementing specific proteins, as indicated – most notably, recombinant human albumin (expressed in *P. pastoris*; Millipore Sigma). Immunodepletion experiments were performed by incubating PPP with Dynabeads coated with indicated antibodies (**Table S2**) according to the manufacturer’s instructions. Immunodepletion to ≤10% of unmodified PPP was confirmed by immunoblot prior to further testing. Spiking experiments were performed by supplementing PPP with indicated proteins (**Table S2**) in amounts adequate to replete plasma to 100-200% of rat PNP values prior to clot formation. FXIII activity was inhibited by the addition of 20 µM 1,3,4,5-Tetramethyl-2-[(2-oxopropyl)thio]imidazolium chloride (T101, Zedira BmbH, Germany) prior to clot formation.

### Fibrin Clot Formation Kinetics and Network Density

Plasma clot formation kinetics were determined in a turbidimetric assay at absorbance 405 nm, as described previously [43-45]. Basal turbidity varied between samples, likely due to varying lipid levels, so baseline turbidity was normalized to zero for all samples. Fibrin network density was assessed by laser scanning confocal microscopy using fluorescently-labeled fibrinogen as a tracer, as described previously [43, 46]. In brief, clots were formed *in vitro* using PPP diluted 1:1 in HBS, in the presence of AlexaFluor488-conjugated fibrinogen (160 µM), phospholipids and tissue factor (final concentrations ≈ 1uM and 0.5pM, respectively; RC Low reagent), and calcium chloride (10 mM). In some experiments, recombinant human albumin was added to the PPP prior to clot formation. Clots were then assessed by laser scanning confocal microscopy followed by ImageJ analysis of the following variables: (1) Fibrin Area: AlexaFluor488-positive integrated optical density (IOD), (2) Fibrin Density: percentage of randomly placed grid crosshairs with intersecting fibrin fibers, as previously reported [44], and (3) Fibrin Porosity: AlexaFluor488-negative IOD.

### Enzyme-Linked Immunosorbent Assays, Immunoblots, Antibodies, and Other Reagents

ELISA assays were used to determine rat PPP concentrations of fibrinolysis pathway components as specified in **Table S2**, all ELISA kits were validated for use with the respective rat proteins and performed according to the manufacturer’s instructions. Immunoblot and magnetic bead pulldown assays were performed using rat-specific antibodies against proteins of interest (**Table S2**). For fibrin clot immunoblot experiments, rat PPP was clotted by addition of 20 nM thrombin and 10 mM calcium chloride at RT for 60 min. The resulting clot was then washed once with PBS and solubilized in M-PER (mammalian-protein extraction reagent) containing protease and phosphatase inhibitor cocktails (**Table S2**). Equal amounts (10 µg) of total protein (mixed with Laemmli buffer with β-mercaptoethanol) were loaded in each lane, resolved by 12% SDS-PAGE, and transferred to a PVDF membrane. The membranes were blocked with 5% nonfat dry milk solution, incubated overnight with respective primary antibodies (see **Table S2** for concentrations) followed by incubation with their corresponding secondary HRP-conjugated antibodies. Semi-quantitative determination of each protein was performed by autoradiography after revealing the antibody-bound protein by enhanced chemiluminescence reaction. The density of the bands on scanned autoradiographs was quantified using ImageJ. Where indicated, the relative quantity of each protein of interest was expressed as a ratio to total fibrin(ogen) within the same clot. See **Table S2** for all other reagents.

### Statistics

One- or two-way ANOVA (analysis of variance) was used for multiple group comparisons. When a significant difference was identified by ANOVA, post hoc tests were performed using the Student–Newman–Keuls technique. GraphPad Prism (Boston, MA) software was used for all statistical analyses and for preparation of figures. Statistical significance was defined as *P*<0.05. Data are presented as mean ± SE.

## RESULTS

### Rotational Thromboelastometry Revealed a Hypofibrinolytic Defect in Hypoalbuminemic Whole Blood

Using a ROTEM assay (INTEM activator reagent with added uPA), we previously observed less degradation of clot firmness over time in PAN-NS whole blood, strongly suggestive of a hypofibrinolytic defect [14]. Here we show that this signal was reproducible in a standard ROTEM assay (INTEM without uPA) in which LY60 was directly proportional to plasma albumin (**Fig. 1A-C**). To confirm that this signal was fibrinolysis-dependent, we performed additional ROTEM analyses without and with the addition of tranexamic acid, a potent fibrinolysis inhibitor. These experiments demonstrated that LY60 was significantly reduced by tranexamic acid independently of albumin concentration (**Fig. 1D-E**). We next sought to validate these observations in the *pDTR*-NS rat model in which hypoalbuminemia, proteinuria, and hypercoagulopathy are proportional to DT dose (**Fig. S1**), consistent with prior observations in PAN-NS [14, 34]. ROTEM revealed decreased LY60 in hypoalbuminemic *pDTR*-NS whole blood (**Fig. 1F-H**). Tranexamic acid significantly reduced LY60 independently of albumin concentration in this model as well (**Fig. 1I-J**). These data confirm that ROTEM LY60 is a fibrinolysis-specific variable and suggest that nephrotic hypoalbuminemia is associated with reduced clot lysis.

**Figure 1:**
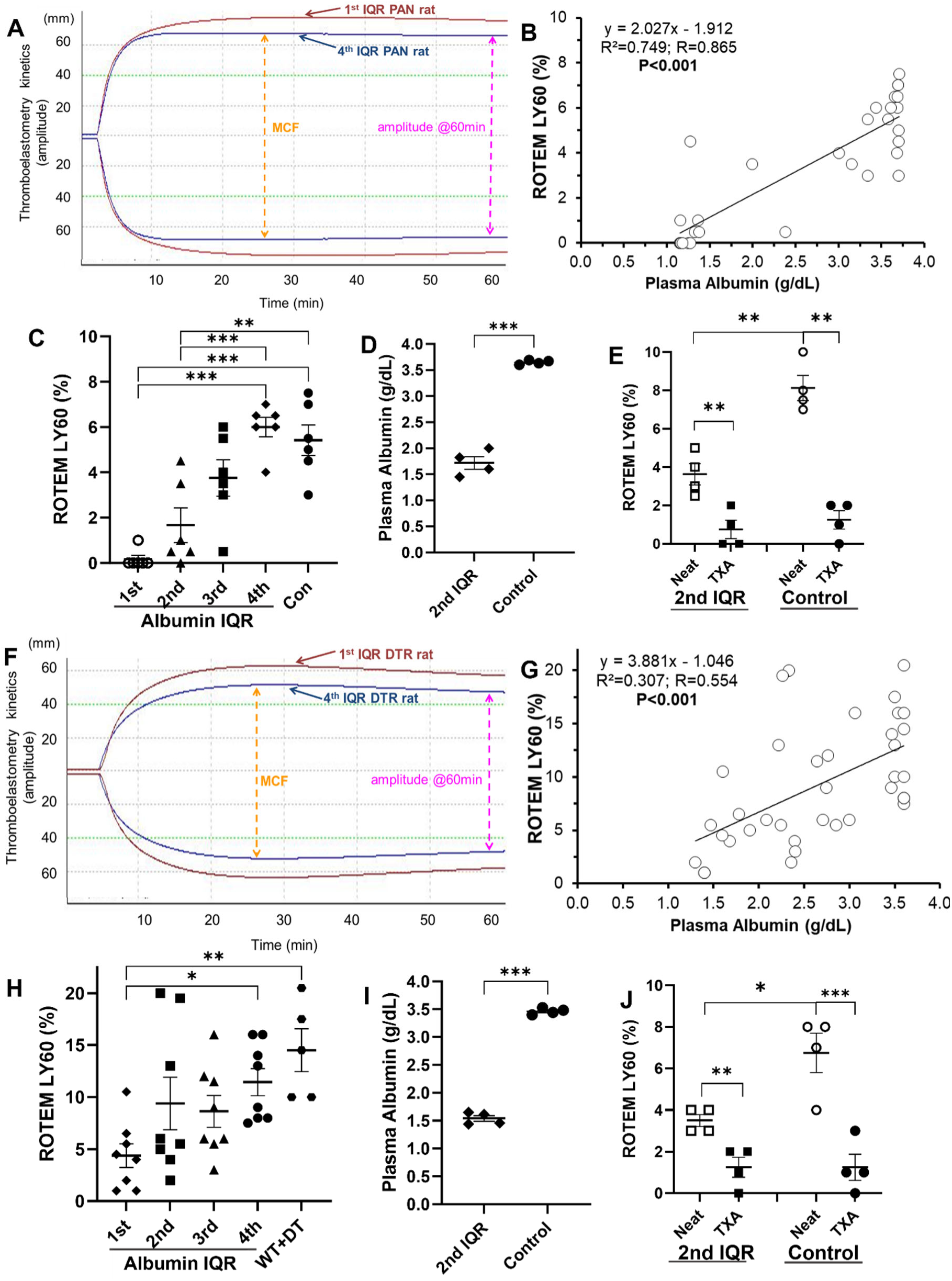
Rotational Thromboelastometry Revealed a Hypofibrinolytic Defect in Hypoalbuminemic Whole Blood. (**A**, **F**) Representative ROTEM tracings from 1^st^ and 4^th^ albumin IQR PAN-(**A**) and *pDTR*-(**F**) nephrotic rats. (**B**, **G**) %Lysis (LY60) determined by ROTEM was directly correlated with plasma albumin concentration in PAN-(**B**) and *pDTR*-(**G**) nephrotic rats. (**C**, **H**) %Lysis by albumin IQR in PAN-(**C**) and *pDTR*-(**H**) nephrotic rats with healthy control rats for both models. (**D**-**E, I**-**J**) Additional groups of PAN- and *pDTR*-nephrotic rats with and without hypoalbuminemia (**D**, **I**) demonstrated that TXA impaired lysis independently of albumin levels (**E**, **J**). ROTEM: rotational thromboelastometry; IQR: interquartile range; PAN: puromycin aminonucleoside; *pDTR*: podocin-promoter driven diphtheria toxin receptor transgene; LY60: Lysis at 60 minutes; TXA: tranexamic acid;

### Hypofibrinolysis was Proportional to Hypoalbuminemia

We next assessed the effect of albumin on clot lysis in PPP. In rats with PAN-NS (**Fig. 2A-B**) or *pDTR*-NS (**Fig. 2C-D**), as well as in NS patient plasma clots (**Fig. 2E-F**), uPA-mediated lysis decreased proportionally to reduced albumin concentrations. A modified CLA performed on *pDTR*-NS PPP clots revealed minimal lysis when uPA was omitted (**Fig. S2A**). Albumin concentration-dependent fibrinolysis persisted when a lower concentration of thrombin (5 nM instead of 20 nM) was used to form the clot (**Fig. S2B**), suggesting that hypofibrinolysis due to hypoalbuminemia is not dependent upon thrombin concentration [14, 34, 36, 45, 47]. In this assay, fibrinolysis is dependent upon uPA-mediated conversion of endogenous plasminogen to plasmin. However, initiation of lysis with both uPA and exogenous plasmin did not meaningfully alter the results (**Fig. S2C**), suggesting that the defect is not due to increased activity of plasminogen activator inhibitors. Quadrupling the uPA concentration did not enable complete lysis of hypoalbuminemic clots despite concomitantly increasing the assay duration to 24 hours (**Fig. S2D**). Performing the assay with tPA (**Fig. S2E**) or plasmin (**Fig. S2F**) in lieu of uPA gave similar results. Collectively, these data suggest that clots formed from hypoalbuminemic plasma are inherently resistant to fibrinolysis.

**Figure 2:**
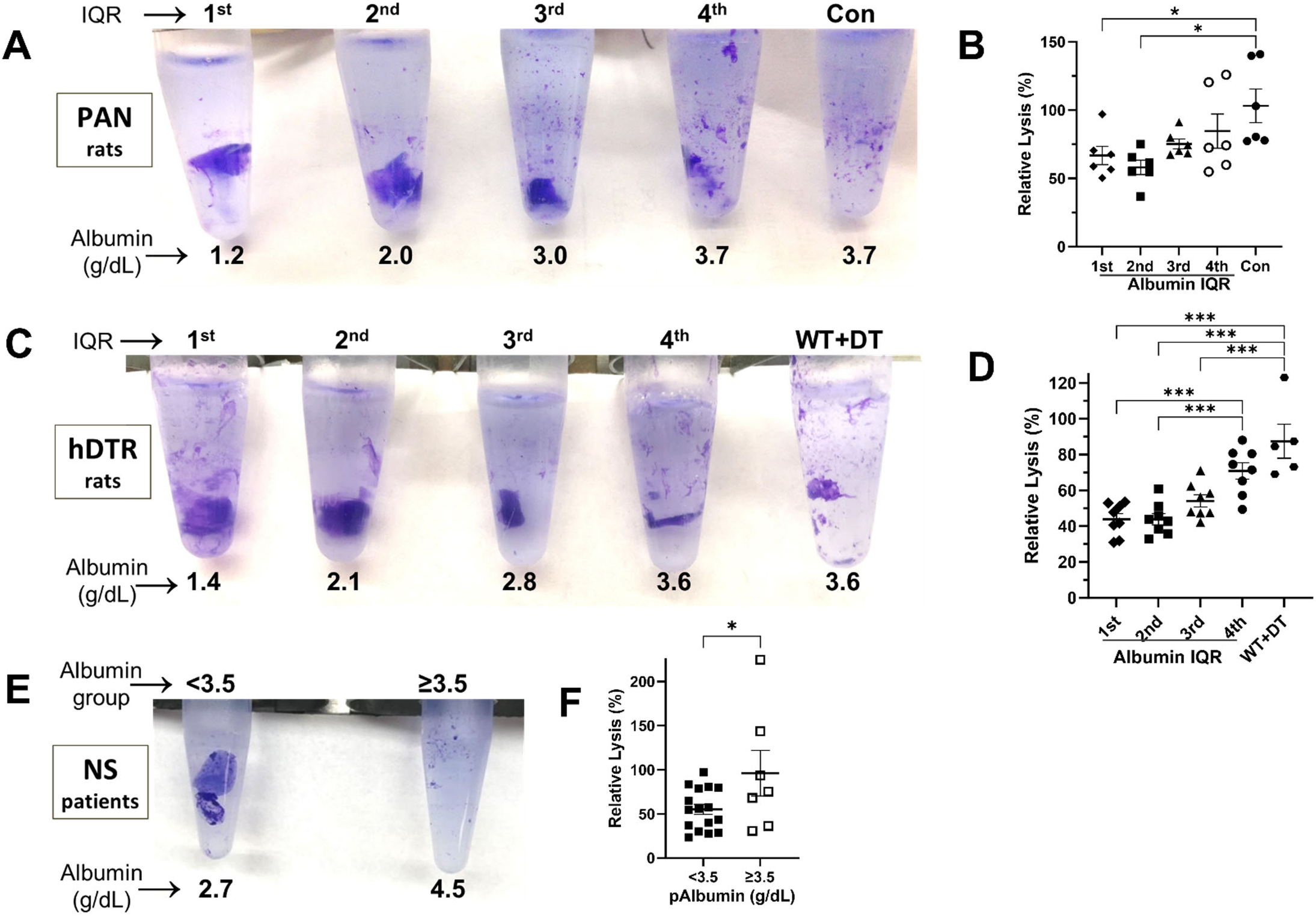
Hypofibrinolysis was Proportional to Hypoalbuminemia. (**A**, **C**, **E**) Representative residual clots at the clot lysis assay endpoint (60 minutes) from PAN- (**A**) or *pDTR*- (**C**) nephrotic rat plasmas and from nephrotic patient plasmas (**E**; annotated with albumin IQR group (**A, C**) or level (**E**)). (**B**, **D**, **F**) Lysis, expressed as a percentage of rat (**B**, **D**) or human (**E**) PNP, in PAN- (**B**) and *pDTR*- (**D**) nephrotic rat plasma clots (and healthy control rat clots) and in clots from nephrotic patients with hypoalbuminemia (<3.5 g/dL) vs. normal albumin levels (≥3.5 g/dL). PAN: puromycin aminonucleoside; *pDTR*: podocin-promoter driven diphtheria toxin receptor transgene; PNP: pooled normal plasma; IQR: interquartile range; WT: wild-type; DT: diphtheria toxin; *n*=6 rats/group (**A-B**); *n*=5-8 rats/group (**C-D**); *n*=7-16 patients/group (**E-F**); \**P*<0.05, \*\*\**P*<0.001.

### Fibrin Clot Formation Kinetics were Enhanced in Hypoalbuminemic Plasma

Accelerated fibrin clot formation and peak turbidity are associated with fibrinolytic resistance [45, 48-55]. Thus, we sought to determine if plasma fibrin clot formation kinetics were altered in hypoalbuminemic *pDTR*-NS rat plasma. Interestingly, both fibrin clot formation rate (Vmax) and peak turbidity increased in proportion to hypoalbuminemia severity (**Fig. 3**). These data suggest that albumin may regulate fibrin clot formation kinetics.

**Figure 3:**
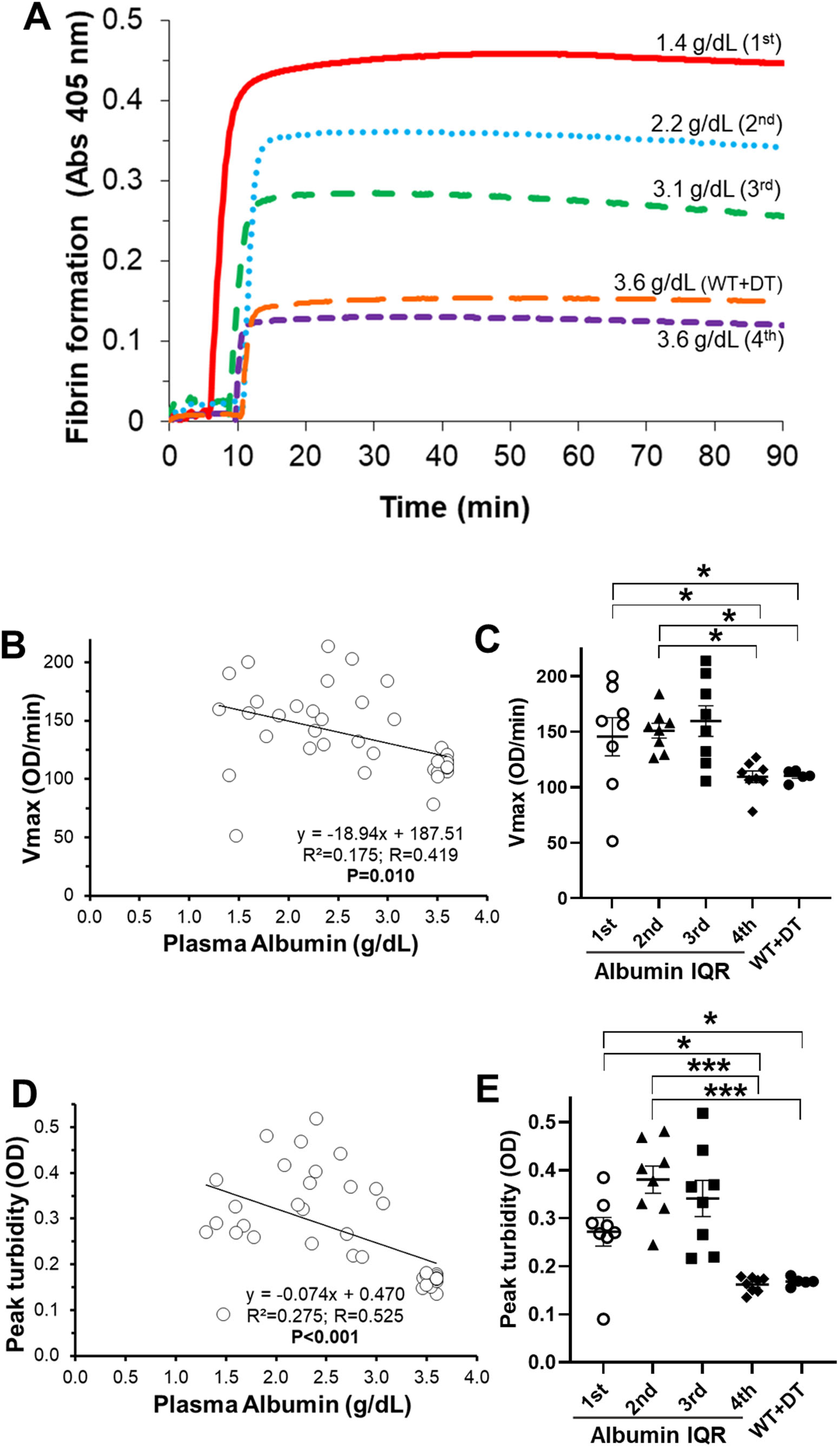
Fibrin Clot Formation Kinetics were Enhanced in Hypoalbuminemic Plasma. (**A**) Representative fibrin clot formation tracings from *pDTR*-nephrotic and a healthy control rat (annotated with plasma albumin levels and albumin IQR group). (**B**) The rate of fibrin formation (Vmax) was inversely proportional to plasma albumin level. (**C**) Vmax by albumin IQR group. (**D**) Peak turbidity was inversely proportional to plasma albumin level. (**E**) Peak turbidity by albumin IQR group. WT: wild-type; DT: diphtheria toxin; *pDTR*: podocin-promoter driven diphtheria toxin receptor transgene; IQR: interquartile range; Vmax: clotting rate; *n*=5-8 rats/group; \**P*<0.05.

### Fibrin Clot Network Density was Inversely Related to Plasma Albumin

Increased thrombin generation, hyperfibrinogenemia, and enhanced plasma clot formation kinetics are co-existent in NS and have all been associated with greater fibrin network density wherein the fibrin fibers are thinner and more tightly packed [14, 44, 45, 56]. Here we found that fibrin area and density increased as albumin concentration decreased in both the *pDTR*-NS rat model and in NS patient plasma (**Fig. 4**) Fibrin network density was significantly and positively correlated with ETP (R=0.498; *P<*0.01), plasma fibrinogen (R=0.146; *P*<0.05), and peak turbidity (R=0.364; *P*<0.05) in the *pDTR*-NS rat model. These data show that fibrin network density is associated with thrombin generation, fibrinogen levels, and fibrin clot formation kinetics during nephrotic syndrome and further demonstrate that density increases during hypoalbuminemia.

**Figure 4:**
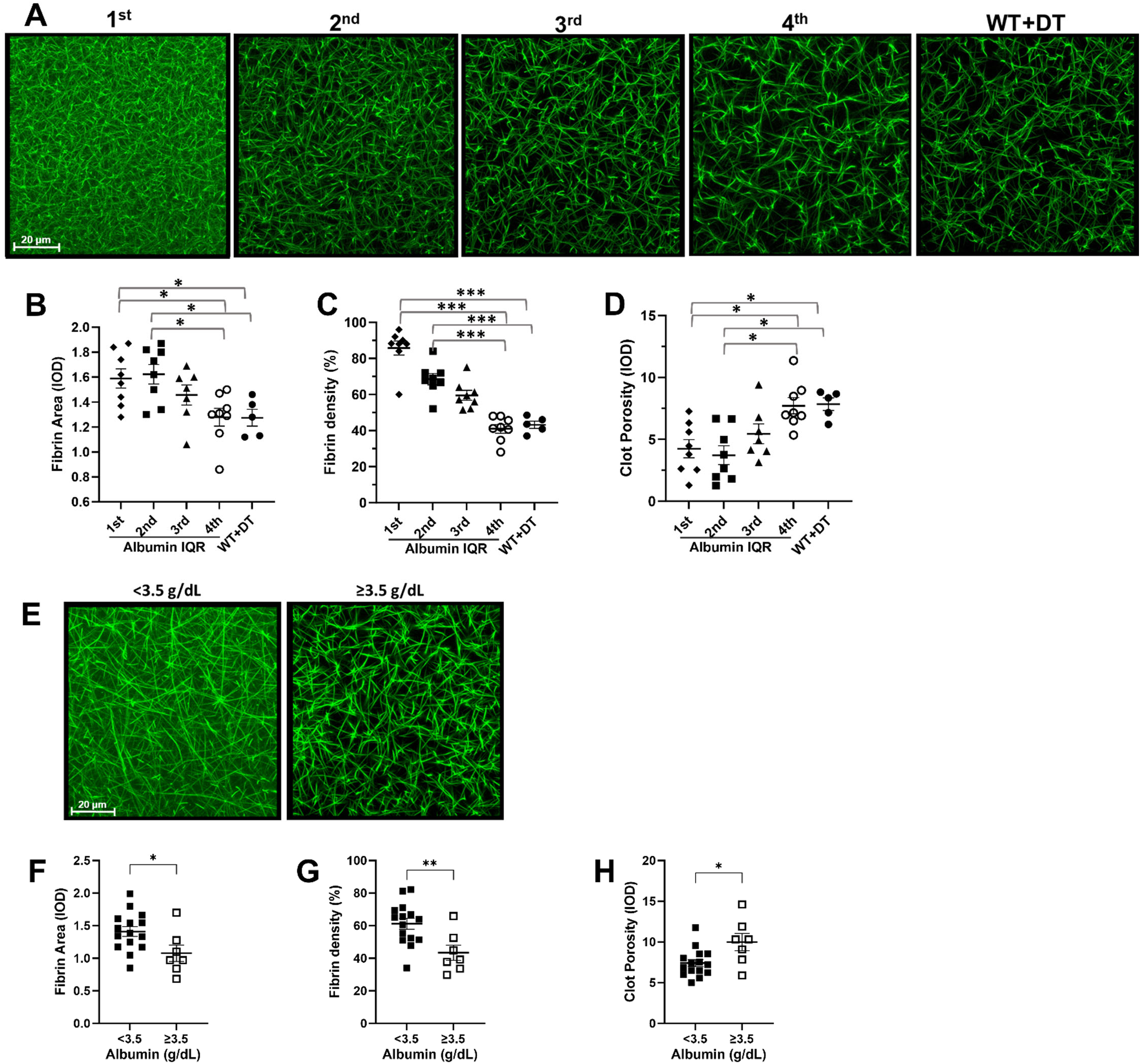
Fibrin Clot Network Density was Inversely Related to Plasma Albumin. Fibrin clots were formed under glass coverslips in the presence of AlexaFlour488-labeled fibrinogen and imaged by laser scanning confocal microscopy. (**A**, **E**) Representative photomicrographs of clots formed from *pDTR*-rat (**A**) and nephrotic patient (**E**) plasma (annotated with albumin IQR group (**A**) or level (**E**)). (**B**, **F**) Fibrin Area (AlexaFlour488-positive integrated optical density (IOD)). (**C**, **G**) Fibrin network density (percentage of randomly placed grid crosshairs with intersecting fibers). (**D**, **H**) Fibrin clot porosity (AlexaFlour488-positive IOD). FITC: fluorescein isothiocyanate; WT: wild-type; DT: diphtheria toxin; *pDTR*: podocin-promoter driven diphtheria toxin receptor transgene; IQR: interquartile range; *n*=5-8 rats/group (**A-E**); *n*=7-16 patients/group (**F-J**); \**P*<0.05, \*\**P*<0.01, \*\*\**P*<0.001.

### Hypoalbuminemic Plasma Clots Contained Less Albumin

To determine the mechanisms mediating altered clot lysis, we quantified the levels of fibrinolysis pathway proteins in PPP. Although both plasminogen deficiency and plasminogen urinary loss have been reported in NS patients [57], plasminogen concentrations were not significantly correlated with albumin in *pDTR*-NS plasma (**Fig. S3A**). Consistent with previous reports, hyperfibrinogenemia was observed in hypoalbuminemic plasma (**Fig. S3B**) [8, 14]. Plasminogen activator inhibitor-1 (PAI-1), α2-macroglobulin, and uPA levels were also significantly increased in hypoalbuminemic plasma (**Fig. S3C-S3E**). There were no significant changes in the concentrations of tPA, TAFI, or α2-AP (**Fig. S3F-S3H**). We next used immunoblotting to determine intra-clot quantities of albumin and fibrinolytic system components in *pDTR*-NS plasma clots relative to plasma albumin levels (**Figs. 5 and S4**). Unsurprisingly, intra-clot albumin was decreased in concert with decreasing plasma albumin concentration (**Fig. 5A**). We expected to find higher quantities of fibrin(ogen) in the densest clots (from the most hypoalbuminemic plasmas). To our surprise, clot fibrin(ogen) content was decreased in hypoalbuminemic plasma (**Fig. 5B**), such that the least dense clots (those with the highest albumin levels) had the highest fibrin(ogen) content. Moreover, the lowest albumin quartile plasma clots had a lower albumin:fibrin(ogen) content ratio than the highest albumin quartile clots (**Fig. 5C**). Interestingly, although the levels of the fibrinolysis inhibitors PAI-1, α2AP, α2M, and TAFI increased with hypoalbuminemia (relative to fibrin(ogen), **Fig. S5**), immunodepleting these inhibitors from nephrotic, hypoalbuminemic plasma did not correct the hypofibrinolytic defect in CLA experiments (**Fig. S6**). Likewise, although intra-clot levels of tPA, uPA, PAI-2, factor XII, factor XI, and prekallikrein decreased with hypoalbuminemia (relative to fibrin(ogen), **Fig. S7**), supplementing plasma to achieve increased levels of these proteins (100-200% of rat PNP) did not correct the lysis in CLA experiments (**Fig. S8**). We also used a transglutaminase inhibitor, T101, to determine if albumin inhibits factor XIII-dependent fibrin cross-linking and thereby enhances fibrinolysis; however, factor XIII inhibition did not significantly alter CLA results (**Fig. S9**). These data show that plasma albumin concentrations may influence the incorporation of various fibrinolytic components into fibrin clots, but that these changes do not directly influence fibrinolytic susceptibility.

**Figure 5:**
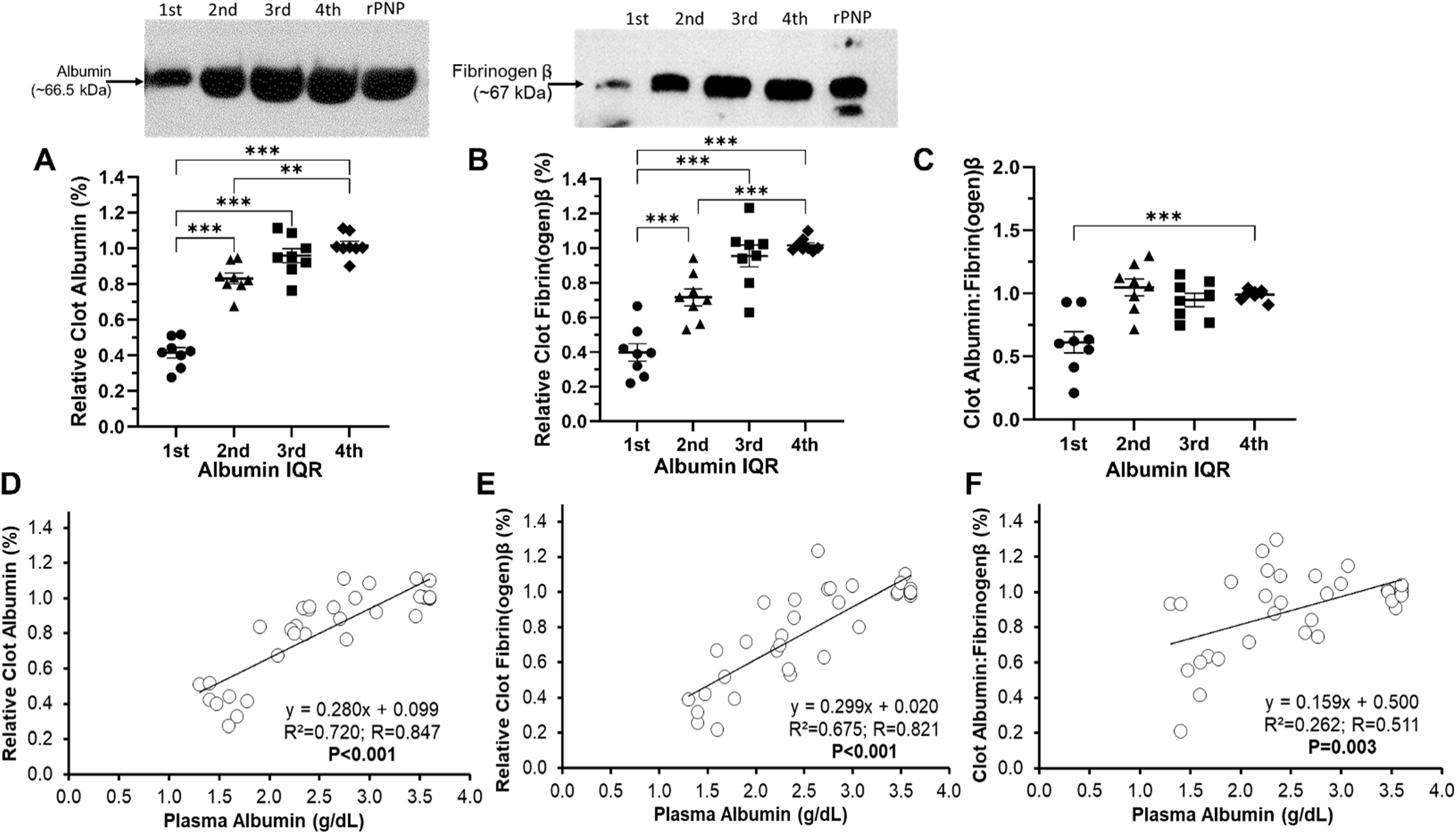
Hypoalbuminemic Plasma Clots Contained Less Albumin. (**A**, **B**) Fibrin clots were formed from *pDTR*-rat plasma, washed, solubilized in M-PER, then immunoblotted for albumin (**A**) and fibrin(ogen) β-chain (**B**); representative immunoblots are shown above each graph. Data are expressed relative to fibrin clots formed from rat PNP (rPNP). (**C**) Intra-clot albumin content (relative to fibrin(ogen)). (**D-F**) Clot content of albumin (**D**) and fibrin(ogen) β-chain (**E**), as well as albumin:fibrin(ogen) ratio (**F**) were directly correlated with plasma albumin. *pDTR*: podocin-promoter driven diphtheria toxin receptor transgene; M-PER: mammalian protein extraction reagent; IQR: interquartile range; n=5-8 rats/group; \*\**P*<0.01, \*\*\**P*<0.001.

### Albumin Repletion Corrected Hypofibrinolysis and Fibrin Network Density

The negative findings with manipulation of fibrinolytic system components prompted us to directly test albumin repletion. Interestingly, dissolving lyophilized recombinant albumin into *pDTR*-NS plasma samples to achieve the upper limit of the healthy rat reference range (4.5 g/dL; 677 µM) enhanced fibrinolysis to values that were no longer significantly different from controls (**Fig. 6**) [58]. In contrast, supplementation with an equimolar amount of a control protein from the serpin protein family (ovalbumin [OvA]) had no effect on fibrinolysis [59, 60]. Moreover, albumin repletion decreased fibrin network density to values no different than control clots following albumin repletion whereas OvA addition had no significant effect (**Fig. 7**). Taken together with the intra-clot albumin data, these observations suggest that clot-incorporated albumin reduces fibrin network density and increases fibrinolytic susceptibility.

**Figure 6:**
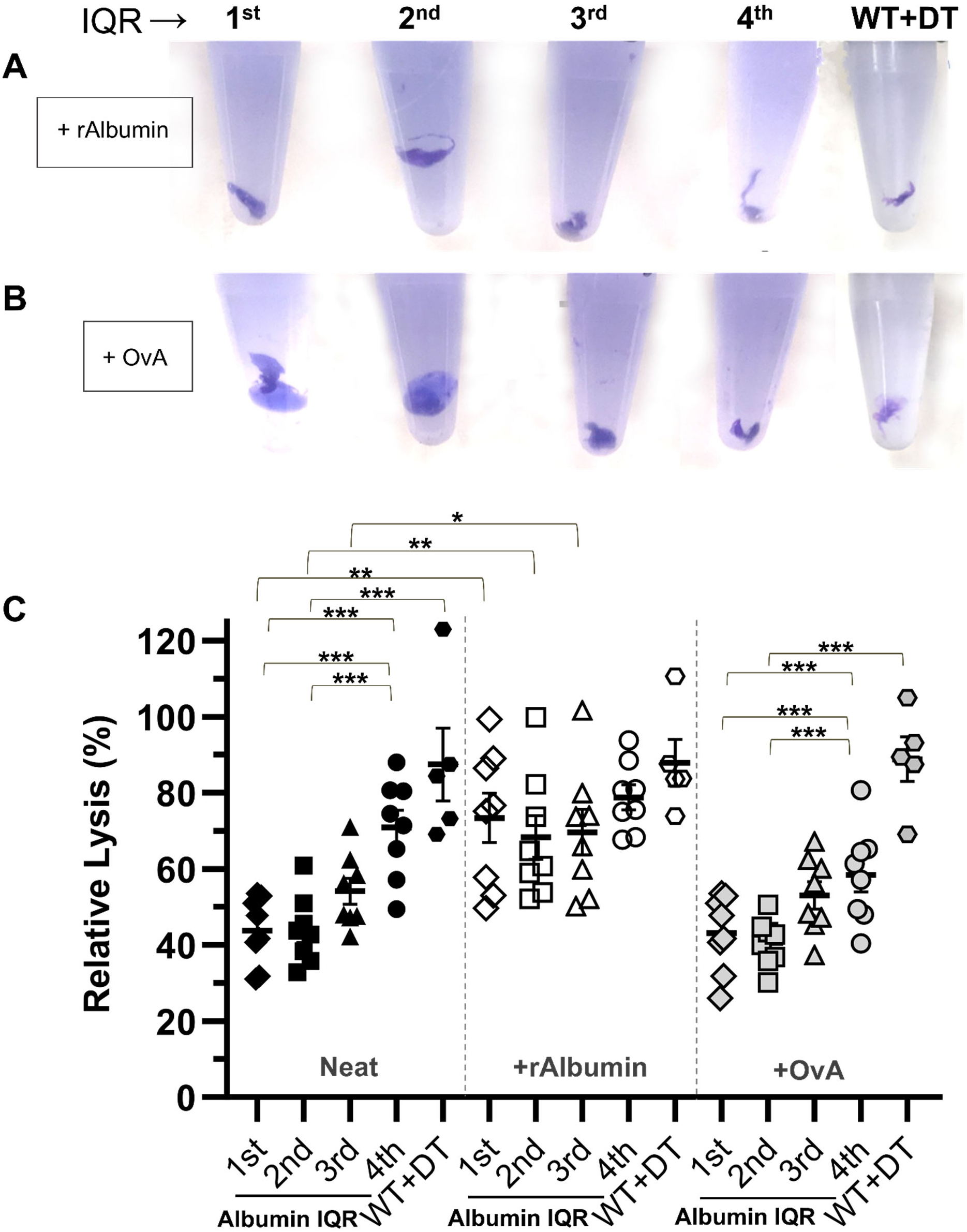
Albumin Repletion Corrected Hypofibrinolysis. (**A**, **B**) Representative residual clots at the clot lysis assay endpoint (60 minutes) from *pDTR*-nephrotic rat plasmas supplemented with recombinant Albumin (**A**) or Ovalbumin (**B**) (Figure 2C shows representative residual clots of unmanipulated samples). (**C**) Lysis, expressed as a percentage of rat PNP values, is corrected by rAlbumin, but not Ovalbumin, supplementation. WT: wild-type; DT: diphtheria toxin; *pDTR*: podocin-promoter driven diphtheria toxin receptor transgene; rAlbumin: recombinant albumin; OvA: ovalbumin (note: OvA is a serpin-family, not an albumin-family, protein); n=5-8 rats/group; \**P*<0.05, \*\**P*<0.01, \*\*\**P*<0.001.

**Figure 7:**
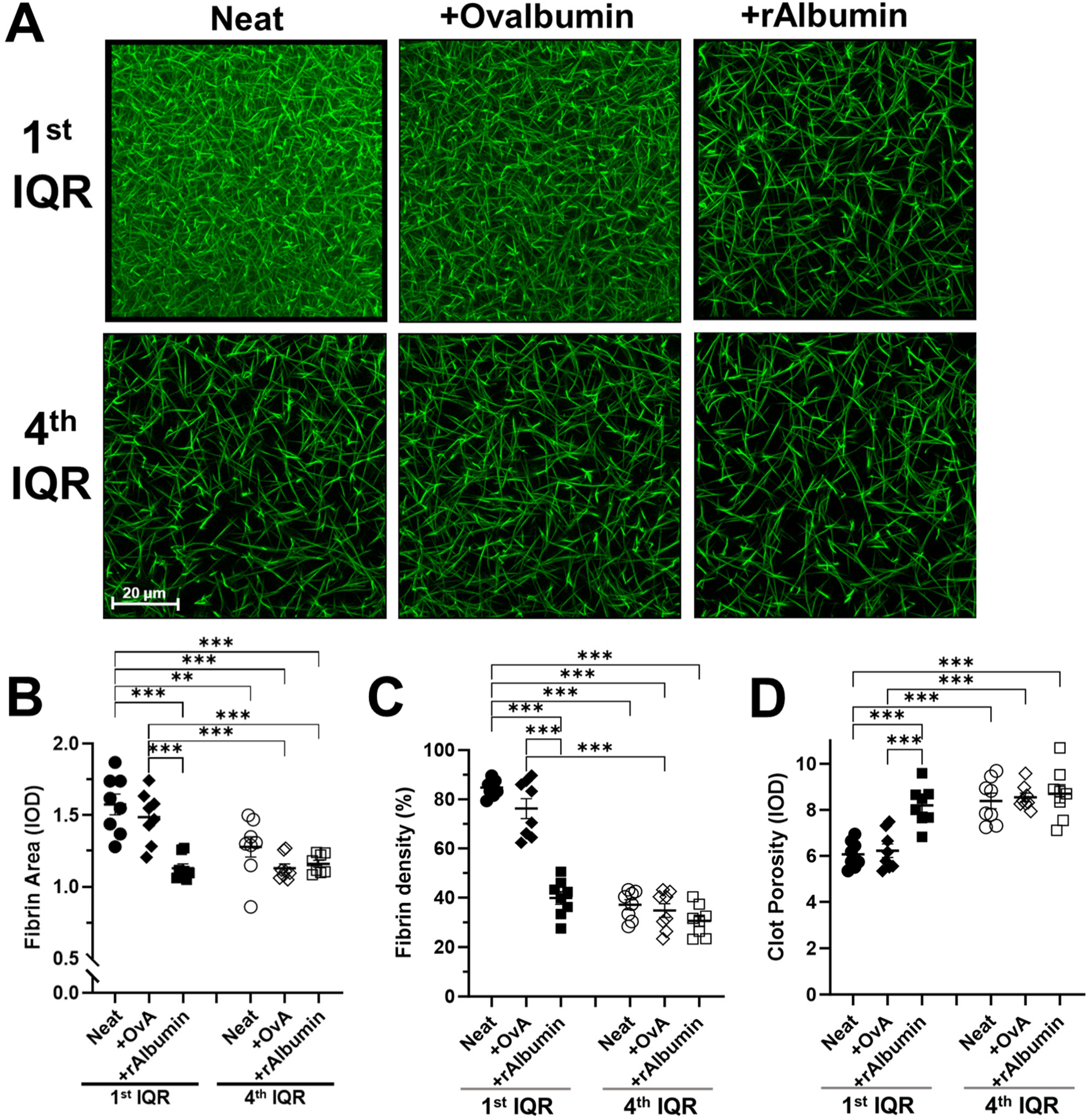
Albumin Repletion Corrected Fibrin Network Density. (**A**) Representative photomicrographs of clots formed from 1^st^ and 4^th^ albumin IQR *pDTR*-rat plasma (as in Figure 4A) with Ovalbumin or rAlbumin supplementation. (**B**) Fibrin area. (**C**) Fibrin network density. (**D**) Fibrin clot porosity. IQR: interquartile range; WT: wild-type; DT: diphtheria toxin; *pDTR*: podocin-promoter driven diphtheria toxin receptor transgene; rAlbumin: recombinant human albumin; OvA: ovalbumin (note: OvA is a serpin-family, not an albumin-family, protein); n=8 rats/group; \**P*<0.05, \*\**P*<0.01, \*\*\**P*<0.001.

### Albumin Dictated Fibrinolytic Susceptibility and Fibrin Network Density in Analbuminemic Rat Plasma

To determine if albumin modulates fibrinolysis in settings beyond nephrotic syndrome, we examined plasma samples from a mutant analbuminemic rat model [29-32]. Consistent with the nephrotic-hypoalbuminemic data presented above, *Alb*^-/-^ plasma clots were resistant to fibrinolysis and this hypofibrinolytic defect was normalized with the addition of albumin (**Fig. 8A**). *Alb*^-/-^ rat blood was also hypofibrinolytic by ROTEM and albumin repletion significantly corrected the LY60 values to that of wild-type (*Alb*^+/+^) controls (**Fig. S10**). Albumin supplementation supported fibrinolysis with similar half-maximal effective concentration (EC_50_) values in both nephrotic and *Alb*^-/-^ plasma (**Fig. S11**). Moreover, fibrin network density was increased in clotted *Alb*^-/-^ plasma and reverted to normal following albumin supplementation (**Fig. 8B**). Collectively, these data confirm that albumin levels are an independent determinant of fibrinolytic susceptibility and fibrin clot density.

**Figure 8:**
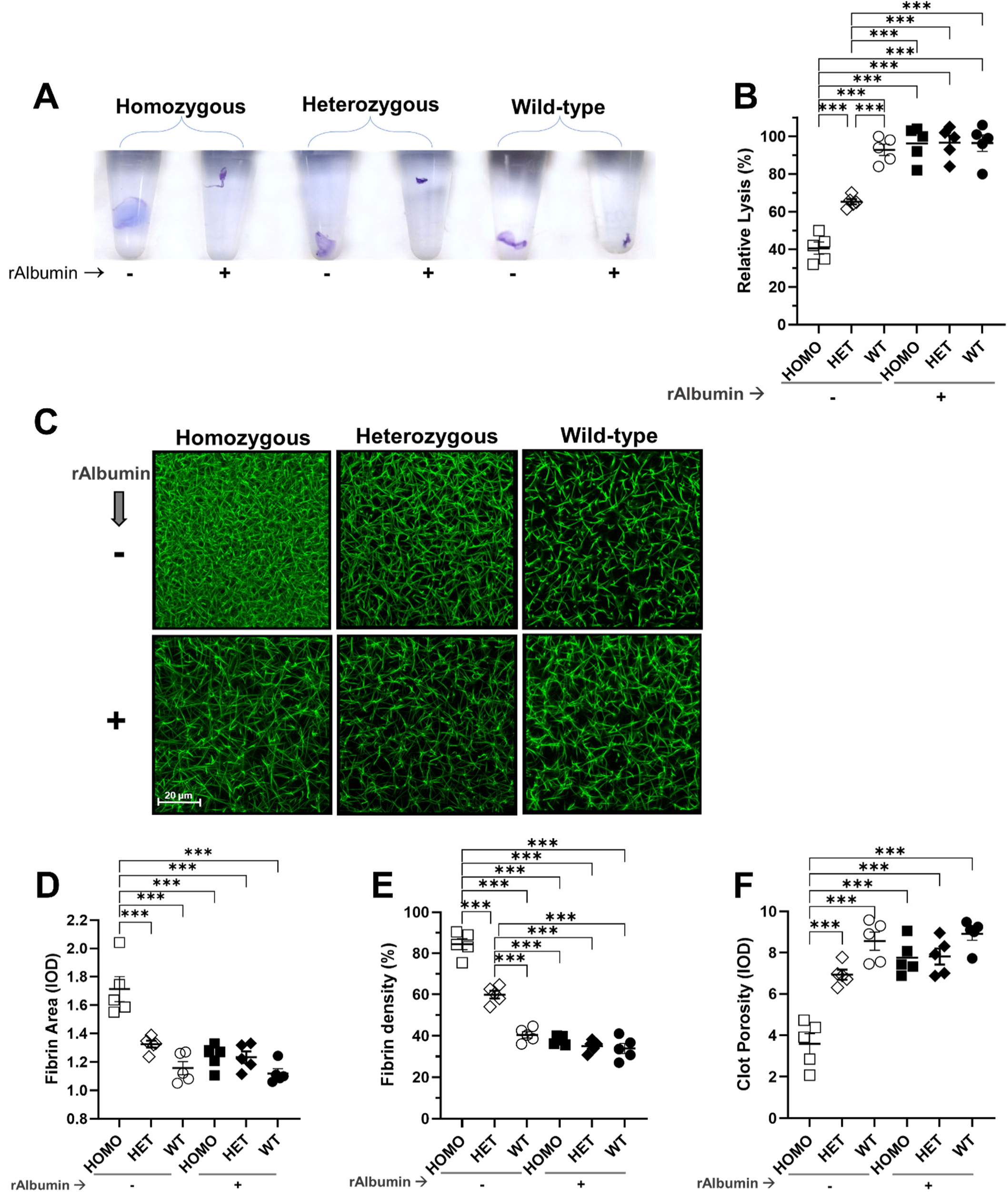
Albumin Dictated Fibrinolytic Susceptibility and Fibrin Network Density in Albumin Knockout Plasma. (**A**) Representative residual clots at the clot lysis assay endpoint (60 minutes) from albumin knockout rat plasmas (as in Figure 4A) without and with recombinant Albumin (rAlbumin) supplementation. (**B**) Lysis, expressed as a percentage of rat PNP values, was diminished in homozygous and heterozygous albumin knockout rat clots and was corrected by rAlbumin supplementation. (**C**) Representative photomicrographs of clots formed from albumin knockout rat plasmas without and with rAlbumin supplementation. (**D**) Fibrin area. (**E**) Fibrin network density. (**F**) Fibrin clot porosity. rAlbumin: recombinant albumin; n=5 rats/group; \*\*\**P*<0.001.

## DISCUSSION

In this study, hypoalbuminemia resulted in dense fibrin clots that were resistant to fibrinolysis and both effects were corrected by restoration of albumin concentrations to healthy control levels. These effects were observed in 2 hypoalbuminemic rat nephrotic syndrome (NS) models, hypoalbuminemic NS patient plasma, and in mutant analbuminemic rats. Importantly, systemic inflammation is not a feature of these rat models, enabling us to discern the effects of hypoalbuminemia independently of the negative acute phase response behavior of albumin that may be related to thromboinflammation [3, 27-32]. Subjecting the clots to various profibrinolytic conditions and immunodepleting fibrinolysis inhibitors did not substantially improve fibrinolysis, suggesting that hypoalbuminemic plasma clots are inherently resistant to fibrinolysis. Multiple studies have demonstrated that dense fibrin clots are less susceptible to fibrinolysis [53, 61-68]. In the present study, hypoalbuminemic clots displayed increased density that was correctable with recombinant albumin supplementation. Together, these data strongly suggest that fibrin clot density and fibrinolytic susceptibility are, at least partly, dictated by plasma albumin levels. Thus, the loss of albumin-dependent effects on fibrin clot structure may help explain why hypoalbuminemic patients are at higher risk for thrombosis and cardiovascular disease [1-4].

We and others have demonstrated hypofibrinolysis by a variety of methods in NS patients and animal models [10-15]. In a previous study, *in vivo* thrombosis modeling in analbuminemic (*Alb*^-/-^) rats revealed larger thrombi and less frequent thrombus embolization compared to albumin sufficient control rats [69]. The authors of that paper speculated that decreased thrombus embolization resulted from diminished *in vivo* fibrinolytic activity. In the present study, we observed diminished fibrinolysis in our *ex vivo* assays with both hypoalbuminemic NS and *Alb*^-/-^ plasmas. The present data strongly suggest that hypoalbuminemia is associated with denser fibrin clots that are resistant to fibrinolysis.

Previous studies have demonstrated that albumin supplementation enhances fibrinolysis of clotted nephrotic syndrome plasma [11, 19]. Our fibrinolysis assay results confirm these observations and further demonstrate the same effect in *Alb*^-/-^ plasma. The latter data provide the most conclusive evidence to date that albumin directly enhances fibrinolysis and are consistent with the data demonstrating decreased thrombus embolization in *Alb*^-/-^ rats [69]. Significantly lower tPA and higher α2-AP activities were observed in *Alb*^-/-^ plasma and, in a purified biochemistry system, both rat and human albumin enhanced tPA activation of plasminogen, leading the authors to conclude that albumin contributes to fibrinolytic activity [69]. Meanwhile, we were unable to demonstrate albumin-dependent enhancement of tPA activity against its chromogenic substrate (data not shown). Nonetheless, the present and prior data together strongly suggest that albumin enhances fibrinolytic capacity. While the molecular mechanism underlying this behavior remains unresolved, the present data suggest that albumin directly influences fibrin clot structure which, in turn, dictates its fibrinolytic susceptibility.

Our results demonstrate that albumin regulates fibrin clot structure resulting in decreased fibrin network density. These data are consistent with previous reports that used a variety of approaches to show that albumin concentrations alter fibrin structure [11, 20, 22, 23]. Under scanning confocal laser microscopy, both nephrotic and *Alb*^-/-^ samples demonstrated increased fibrin network density that was correctable with albumin supplementation, resulting in thicker fibers and increased porosity [11]. In purified systems, at very low concentrations (≤5 μM) albumin decreased turbidity but as the levels increased (6-100 μM) turbidity increased, suggesting that albumin influences fibrin fiber density in a concentration-dependent manner [20]. Unfortunately, those experiments were conducted at non-physiologic fibrin(ogen)-to-albumin molar ratios making the results difficult to put into physiologic context. Interestingly, when fibrin fiber density is estimated by turbidity, albumin has been reported to decrease fiber thickness in purified fibrinogen clots but to increase thickness in clotted plasma samples [22]. Similarly, in magnetic birefringence assays of clotted plasma, albumin appears to enhance fibrin fiber thickness [23]. These latter data suggest that interactions with other plasma components may be involved in the mechanism by which albumin regulates fibrin structure. Collectively, the present and previously published data strongly suggest that, at physiologically relevant concentrations, albumin increases fibrin fiber thickness, decreases network density, and increases porosity. Some authors have proposed volume exclusion as a possible explanation for the latter observations [23]. However, both our data demonstrating that a control protein does not alter fibrin clot structure and the purified fibrinogen reports (which also included control proteins) argue against this possibility [22]. Thus, the molecular mechanism underlying the effect of albumin on fibrin network density remains unexplained.

Whereas the previously published *Alb*^-/-^ rat thrombosis model study is consistent with our *ex vivo* observations, future studies should explore these phenomena using *in vivo* models to more fully understand their physiologic relevance [69]. With the exception of the ROTEM experiments, the present data was predominantly generated in platelet poor plasma, negating potential interactions with cellular blood components and endothelium, emphasizing the need for *in vivo* translation of these observations [70]. Hyperfibrinogenemia is a feature of both NS and the *Alb*^-/-^ rat and hyperfibrinogenemia is associated with both increase clot density and fibrinolytic resistance [8, 14, 44, 69]. While it is likely that hyperfibrinogenemia contributed to the hypofibrinolytic phenotypes shown in these data, albumin was able to modify lysis of these samples, strongly suggesting that albumin has a fibrinogen-independent effect on fibrinolysis. Interestingly, fibrinogen-to-albumin ratio determination has recently emerged as a novel biomarker for thromboembolic risk and outcomes [71, 72]. It remains unclear from the presently available data if albumin regulates clot structure via direct mechanisms (e.g. intramolecular interactions with fibrin(ogen)) or if these effects are indirect (perhaps via interactions with other components of the fibrin clot or coagulation system) [73]. Despite these limitations, the available data strongly suggest that albumin is an important regulator of fibrin clot structure and fibrinolytic susceptibility.

Hypoalbuminemia is associated with both venous and atherothrombotic cardiovascular disease [1-4]. In this study we found that hypofibrinolysis is a feature of hypoalbuminemia. In turn, hypofibrinolysis is associated with increased risk for both venous and arterial thrombosis, suggesting a mechanistic link between these risk factors [16, 17]. Future studies should thus focus on determining the molecular mechanisms by which albumin regulates fibrin clot structure and fibrinolytic susceptibility.

## ACKNOWLEDGEMENTS

This work was supported by grants from the National Institute Health (NIH) grants R01 DK124549 to B. A. Kerlin and R01 HL126974 and R01 HL147894 to A. S. Wolberg as well as a research endowment from the George and Elizabeth Kelly Foundation (Lewis Center, OH) to B. A. Kerlin. The content is solely the responsibility of the authors and does not necessarily represent the official views of the National Institutes of Health. The authors are indebted to Samir V. Parikh, Brad H. Rovin, and William E. Smoyer for recruiting patients into the Columbus Nephrotic Syndrome Cohort. *pDTR* transgenic rat breeders were a kind gift from Roger C. Wiggins and Bruce A. Molitoris.

## AUTHOR CONTRIBUTIONS

BAK directed and supervised all experimental design, data acquisition, and analysis, edited the manuscript and figures, and compiled the manuscript for submission. APW and BAK drafted the manuscript together. ASW critically revised the manuscript, provided important intellectual content, and supervised the fibrin clot formation kinetic studies which were performed in her lab by LAH. APW, KJW, and ZSS conducted all other experiments, analyzed data, and prepared figure components. All authors approved the submitted version of the manuscript.

