## Supplemental Materials and Data for "Hypoalbuminemia Increases Fibrin Clot Density and Impairs Fibrinolysis"

**Hypoalbuminemia is Associated with Increased Fibrin Clot Density and Reduced Fibrinolytic Susceptibility**

**Table S1: ARRIVE Guidelines for Reporting *In Vivo* Animal Experiments**

| | **hDTR Transgenic Rats** | **Fischer-344 Rats** | **PAN Rats** | **Congenitally Analbuminemic Rats** |
|---|---|---|---|---|
| **Total Rats** | N=32 hDTR rats; males | N=5 Fischer-344 rats; males | N=30 Wistar rats; males | N=18 NAGase (NAR) rats; males & females |
| **Toxin-dependent disease induction** | Diptheria Toxin (DT); 0-75 ng/kg, in 1 mL sterile saline, IP, once | Diptheria Toxin (DT); 50 ng/kg, in 1 mL sterile saline, IP, once | Puromycin aminonucleoside (PAN); 0-150 mg/kg, in 500 uL sterile saline; IV; once | None |
| **Groups (N/group)** | i) IQRs ($1^{st}$-$3^{rd}$) = DT @ 25-75 ng/kg (N=8/group)<br>ii) $4^{th}$ IQR (healthy controls) = DT @ 0 ng/kg (saline only) (N=8) | i) WT-DT (healthy controls) = DT @ 50 ng/kg. (N=5) | i) IQRs ($1^{st}$-$4^{th}$) = PAN @ 25, 50, 100, & 150 mg/kg. (N=6/group)<br>ii) Controls = PAN @ 0 mg/kg (saline only). (N=6) | i) NAR-/- Analbuminemic (N=6)<br>ii) NAR+/- Heterozygous (N=6)<br>iii) NAR+/+ Wildtype (N=6) |
| **Manuscript** | Figures 1, 2, 3, 4, 5, 6, 7, S1, S2, S3, S4, S5, S6, S7, S8, S9 | Figures 1, 2, 3, 4, 6, S1, S2, S3 | Figures 1, 2 | Figures 8, S10 |
| **Husbandry** | Rats were housed in pairs in NexGen Rat 900 cages (Allentown Inc) with standard bedding, automatic water, and chow (Teklad Irradiated Rodent Diet 2920x) provided ad libitum. Environmental enrichment included nesting material (Crink-l'Nest, The Andersons), shelter (Rat Retreats, Bio-serv) and chew toys (Wood Gnawing Blocks, Bio-serv). The room was on a 12:12 light:dark cycle (6AM:6PM), and temperature and humidity were maintained at 72±2 ºF and 30-60%, respectively. | | | |
| **Monitoring** | Rats were monitored daily for overall health and behavior, including loss of response to mild stimuli, lethargy, body condition score, failure to eat or drink, diarrhea, skin lesions or loss of grooming. Body weight was measured and recorded twice weekly.<br>Humane Endpoint criteria: animals that exhibit extreme lethargy/recumbency, have evidence of signs of distress, uncontrollable bleeding, a Body Condition Score ≤2, or weight loss ≥20% of starting body mass.<br>No unexpected or adverse events occurred during the study. No interventions to reduce pain or distress were anticipated or required. | | | |
| **Notes** | Animals randomized to groups; Investigators performing sample collection blinded | | | |

IP: intraperitoneal, IV: intravenous, hDTR: human diphtheria toxin receptor, PAN: puromycin

**Table S2: Enzyme-Linked Immunosorbent Assays, Immunoblots, Antibodies, and Other Reagents**

| Reagent Name | Company | Company Location |
|---|---|---|
| AlexaFluor488-conjugated fibrinogen | Thermo Fisher Scientific | Waltham, MA |
| tissue-type plasminogen activator (tPA) | Diapharma (Technoclone) | West Chester, OH |
| QuantiChrom BCP | BioAssay Systems | Hayward, CA |
| Tranexamic acid | Thermo Fisher Scientific | Waltham, MA |
| Plasmin (human) | Haematologic Technologies | Essex Junction, VT |
| Coomassie Brilliant Blue R-250 | TCI Chemicals | Portland, OR |
| Phosphate Buffered Saline (PBS); Hyclone | GE Healthcare | Logan, UT |
| Urokinase (UK) | Sigma-Aldrich | St Louis, MO |
| alpha-thrombin (human) | Haematologic Technologies | Essex Junction, VT |
| Rox prothrombin assay | DiaPharma | West Chester, OH |
| Technothrombin TGA kit | DiaPharma | West Chester OH |
| Dynabeads | Thermo-Fisher Scientific | Waltham, MA |
| RC Low reagent | DiaPharma | West Chester, OH |
| Fibrinogen antibody | LS Bio | Shirley, MA |
| Alpha-2 macroglobulin antibody | MyBioSource Inc | San Diego, CA |
| Alpha-2 antiplasmin antibody | Santa Cruz Biotechnology | Dallas, TX |
| PAI-1 antibody | MyBioSource Inc | San Diego, CA |
| TAFI antibody | MyBioSource Inc | San Diego, CA |
| Plasminogen antibody | Santa Cruz Biotechnology | Dallas, TX |
| Albumin antibody | Cell Signaling Technology | Danvers, MA |
| tPA antibody | Bioss Inc | Woburn, MA |
| Urokinase antibody | Santa Cruz Biotechnology | Dallas, TX |
| PAI-2 antibody | MyBiosource Inc | San Diego, CA |
| FXI antibody | MyBiosource Inc | San Diego, CA |
| FXII antibody | Novus Biologicals | Centennial, CO |
| Prekallikrein antibody | MyBiosource Inc | San Diego, CA |
| ROTEM reagents (star-tem, in-tem) | TEM Systems Inc | Durham, NC |
| Rat and Human specific ELISAs | MyBioSource Inc | San Diego, CA |
| Recombinant human albumin from | Sigma-Aldrich | St Louis, MO |
| Ovalbumin | Sigma-Aldrich | St Louis, MO |
| Corn Trypsin Inhibitor | CTI, Prolytix | Essex Junction, VT |
| T101 | Zedira BmbH | Darmstadt,Germany |
| Quantichrom BCP assay kit | Bioassay systems | Hayward, CA |
| Urine protein:creatinine measurement | Antech diagnostics | Morrisville, NC |
| Puromycin aminonucleoside | Millipore Sigma | Burlington, MA |
| Diphtheria Toxin | Sigma-Aldrich | St Louis, MO |
| Prolong gold with DAPI | Invitrogen | Carlsbad, CA |
| Zeiss LSM 700 microscope | Carl Zeiss Microscopy, LLC | White Plains, NY |
| ZEN blue | Zeiss USA | Thornwood, NY |
| Leica DMI 4000B fluorescence microscope | Leica | Deerfield, IL |
| ZEN Black software | Zeiss USA | Thornwood, NY |
| Spectramax M2 fluorescent plate reader | Molecular devices | Sunnyvale, CA |
| GraphPad Prism | GraphPad Software | Boston, MA |

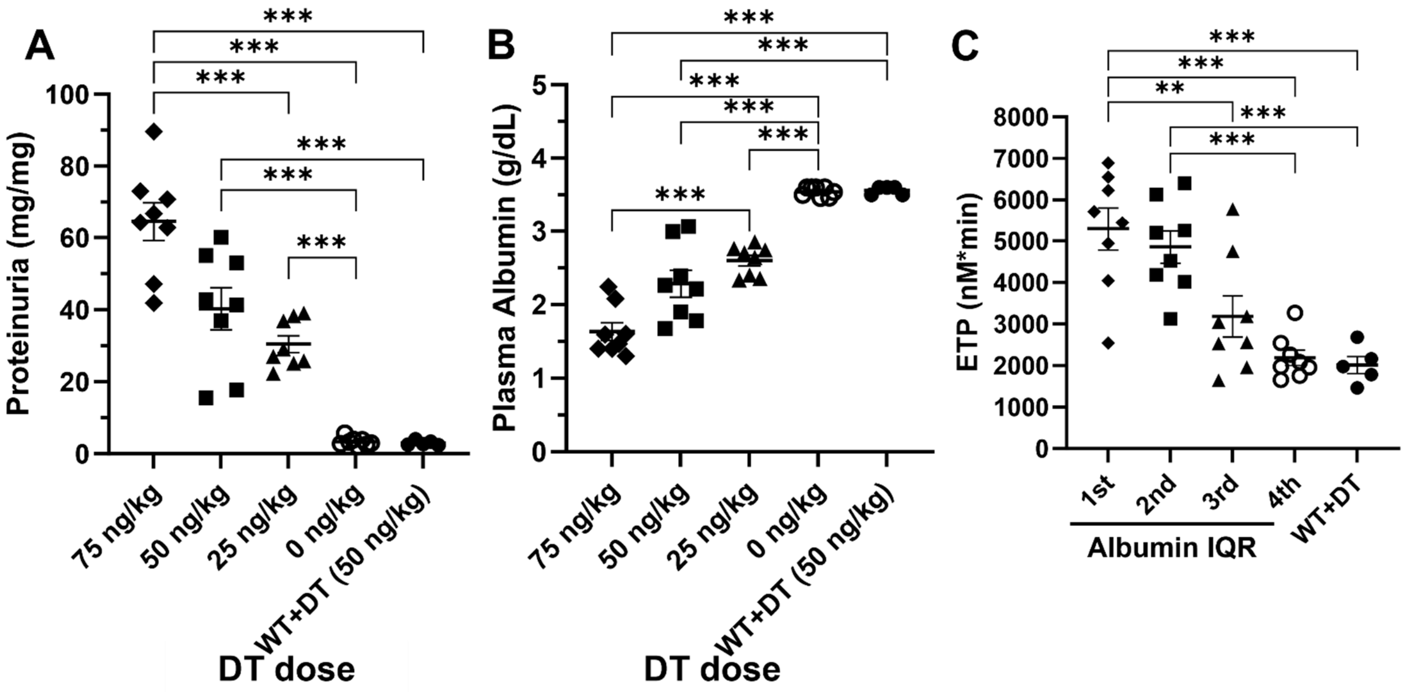


**Figure S1: Dose-Dependent Proteinuria, Hypoalbuminemia, and Hypercoagulopathy in *pDTR* Rats.** (**A**) Proteinuria (expressed as protein (mg) to creatinine (mg) ratio) increases in a diphtheria toxin dose-dependent manner in *pDTR* rats (Note: diphtheria toxin doses are plotted in descending order on x-axis). (**B**) Plasma albumin concentrations decrease in a diphtheria toxin dose-dependent manner in *pDTR* rats (Note: diphtheria toxin doses are plotted in descending order on x-axis). (**C**) Endogenous thrombin potential increases in the setting of hypoalbuminemia in *pDTR* rats. *pDTR*: podocin-promoter driven diphtheria toxin receptor transgene; WT: wild-type; DT: diphtheria toxin; IQR: interquartile range; ETP: endogenous thrombin potential; $n$=5-8 rats/group; **$P$<0.01, ***$P$<0.001.

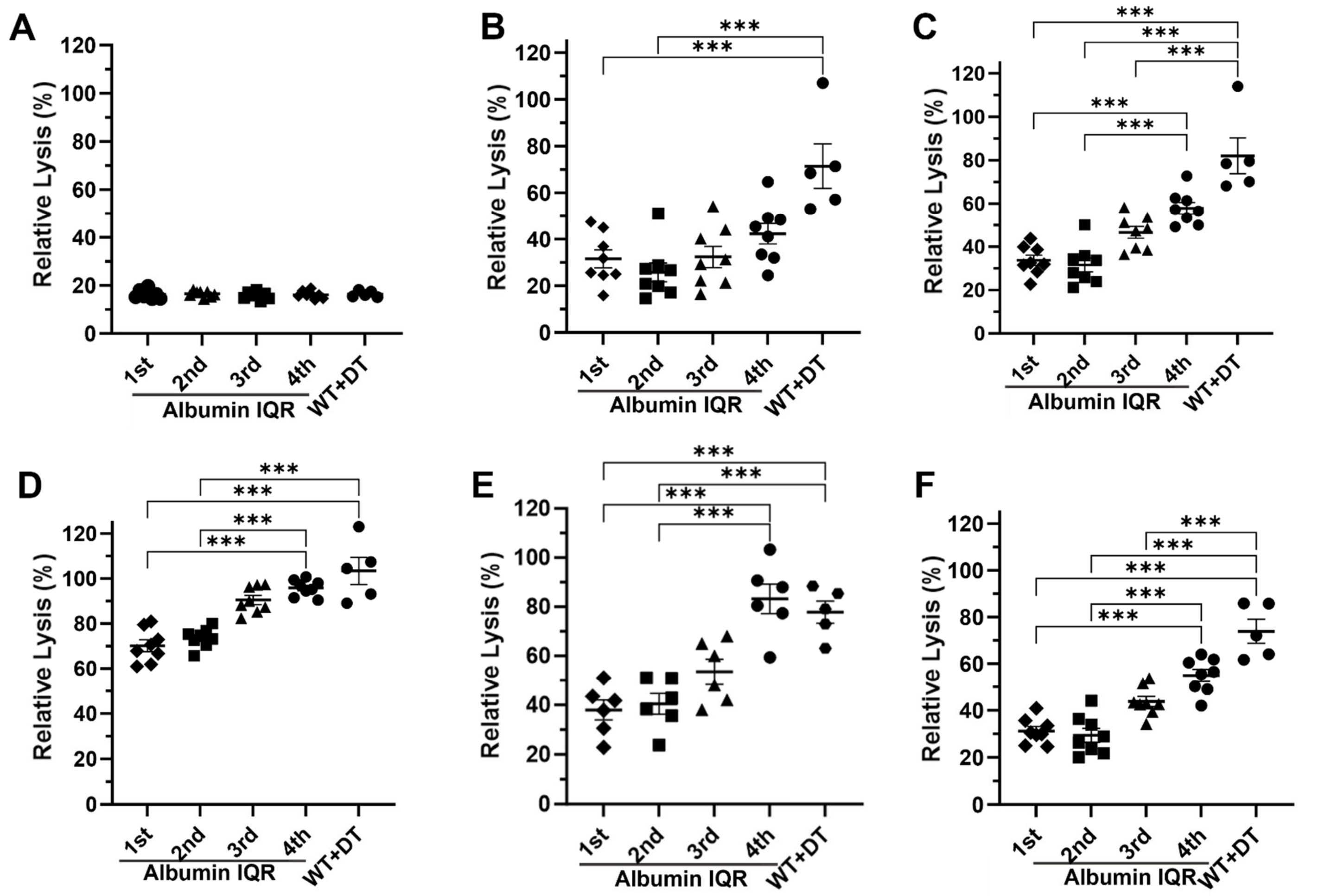


**Figure S2: Clot Lysis is Albumin-Dependent Under Varying Conditions.** (**A**) Lysis of plasma clots formed with thrombin (20 nM) is minimal and not albumin-dependent in the absence of urokinase (0 IU). (**B**) Urokinase (50 IU)-mediated lysis is albumin-dependent when the clots are formed with low-dose thrombin (5 nM). (**C**) Addition of plasmin (1.2 µM) does not alter the albumin-dependent lysis of thrombin (20nM)-formed clots in the presence of urokinase (50 IU). (**D**) Albumin-dependent lysis of thrombin (20nM)-formed clots persists when urokinase concentration is increased to 200 IU and endpoint extended to 24 hours at 37 °C. (**E**) Lysis of thrombin (20nM)-formed clots is albumin-dependent when tissue-type plasminogen activator (20 nM) is used instead of urokinase. (**F**) Lysis of thrombin (20nM)-formed clots is albumin-dependent when plasmin (1.2 µM) is used instead of a plasminogen activator. IQR: interquartile range; WT: wild-type; DT: diphtheria toxin; *n*=5-8 rats/group; ***$P$<0.001.

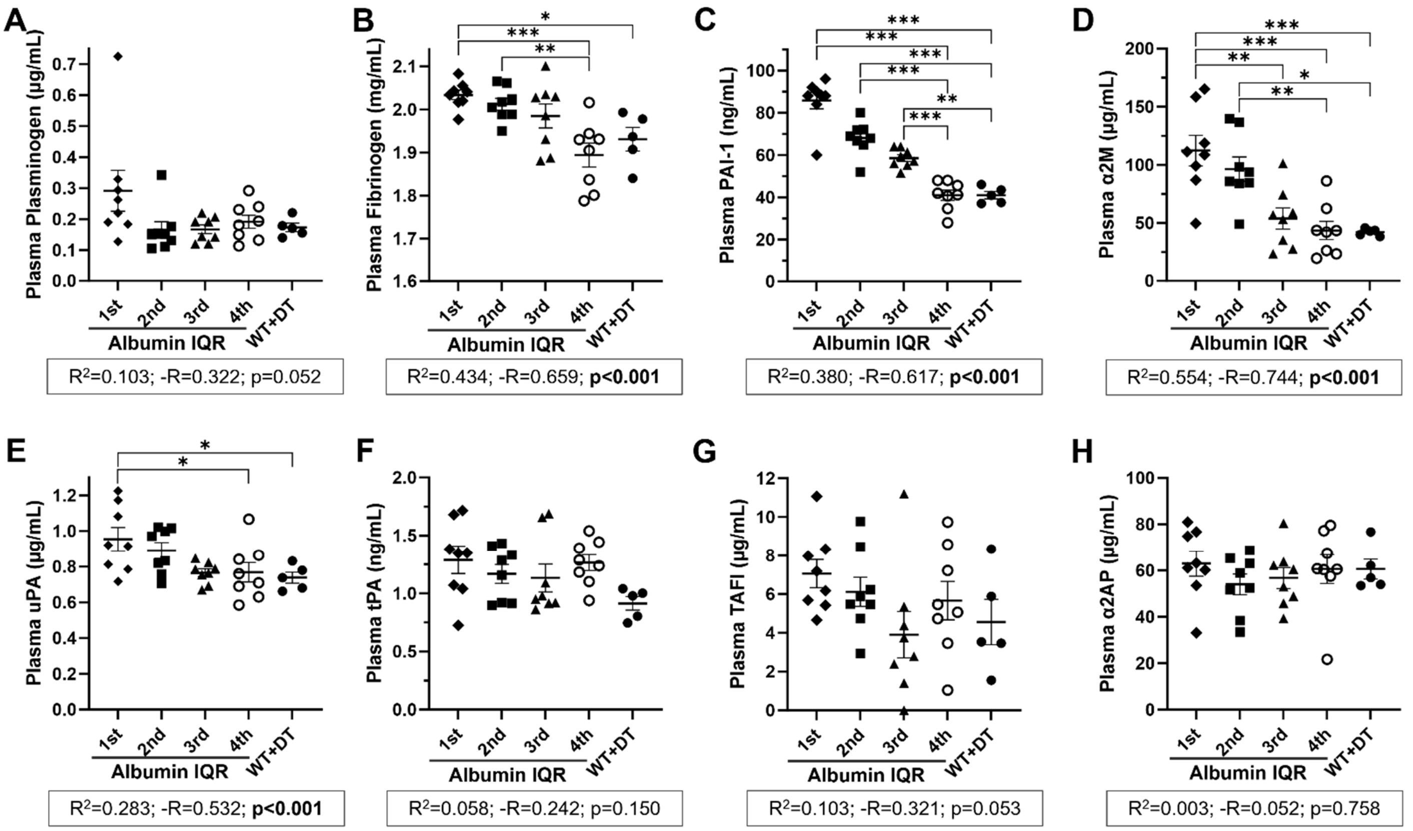


**Figure S3: Fibrinolytic System Proteins in Hypoalbuminemic Nephrotic Plasma.** (**A**) Plasma plasminogen concentrations were not affected by hypoalbuminemia in *pDTR* rat plasma. (**B-E**) Fibrinogen (**B**), PAI-1 (**C**), α2M (**D**), and uPA (**E**) levels increased with decreasing plasma albumin. (**F-H**) tPA (**F**), TAFI (**G**), and α2AP (**H**) levels were unchanged in *pDTR* rat plasma. *pDTR*: podocin-promoter driven diphtheria toxin receptor transgene; IQR: interquartile range; WT: wild-type; DT: diphtheria toxin; PAI-1: plasminogen activator inhibitor-1; α2M: α2-macroglobulin; uPA: urokinase; tPA: tissue-type plasminogen activator; TAFI: thrombin activatable fibrinolysis inhibitor; α2AP: α2-antiplasmin; *n*=5-8 rats/group; *$P$<0.05, **$P$<0.01, ***$P$<0.001.

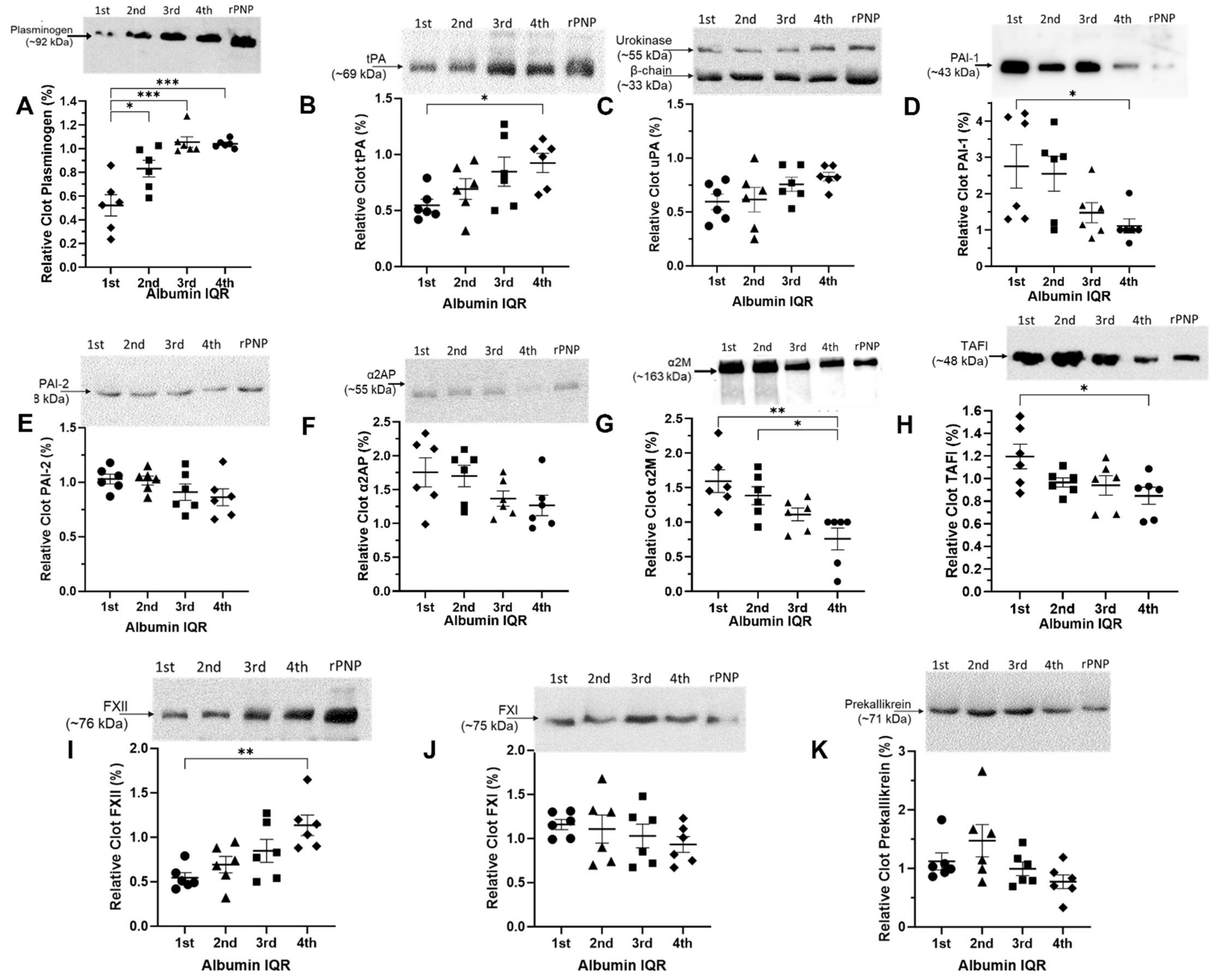


**Figure S4: Fibrinolytic System Protein Incorporation into Clots formed from Hypoalbuminemic Plasma.** Data are expressed relative to fibrin clots formed from rat PNP (rPNP). (**A-C**) Profibrinolytic proteins: Incorporation of plasminogen (**A**) and its activator, tPA (**B**), into clots formed from hypoalbuminemic *pDTR* rat plasma was decreased whereas uPA (**C**) was unchanged. (**D-E**) Plasminogen activator inhibitors: PAI-1 (**D**), but not PAI-2 (**E**), was increased in hypoalbuminemic plasma clots. (**F-H**) Plasmin inhibitors: α2AP (**F**) was unchanged whereas the levels of α2M (**G**) and TAFI (**H**) increased in hypoalbuminemic clots. (**I-K**) Contact pathway enzymes: Factor XII (**I**) incorporation was diminished in hypoalbuminemic clots whereas factor XI (**J**) and prekallikrein (**K**) were unchanged. IQR: interquartile range; tPA: tissue-type plasminogen activator; uPA: urokinase; PAI-1: plasminogen activator inhibitor-1; PAI-2: plasminogen activator inhibitor-2; α2AP: α2-antiplasmin; α2M: α2-macroglobulin; TAFI: thrombin activatable fibrinolysis inhibitor; FXII: factor XII; FXI: factor XI; *pDTR*: podocin-promoter driven diphtheria toxin receptor transgene; *n*=6 rats/group; **P*<0.05, ***P*<0.01, ****P*<0.001.

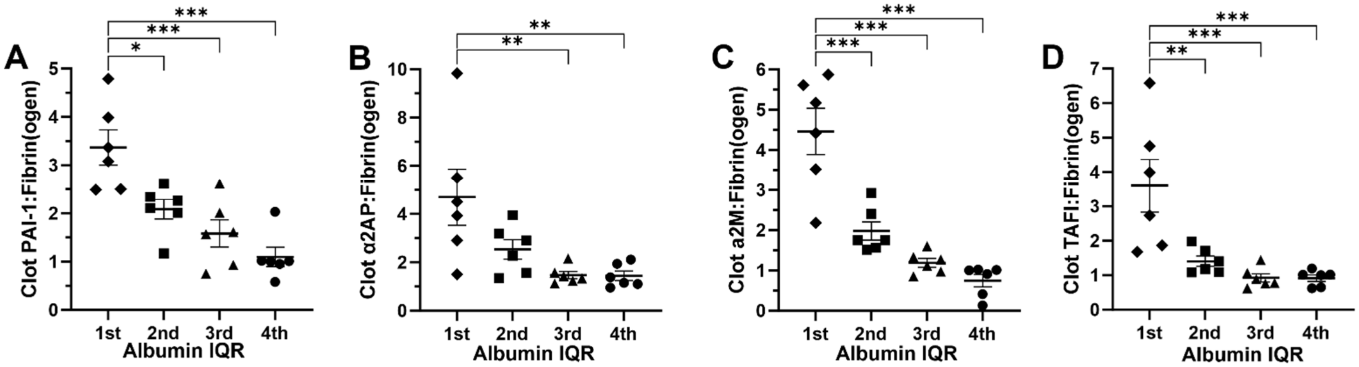


**Figure S5: Fibrinolytic System Proteins that Increased Relative to Fibrin(ogen) in Clots formed from Hypoalbuminemic Plasma.** (**A-D**) Levels of PAI-1 (**A**), α2AP (**B**), α2M (**C**), and TAFI (**D**) were increased relative to fibrin(ogen) in clots formed from hypoalbuminemic *pDTR* rat plasma. PAI-1: plasminogen activator inhibitor-1; α2AP: α2-antiplasmin; α2M: α2-macroglobulin; TAFI: thrombin activatable fibrinolysis inhibitor; IQR: interquartile range; *pDTR*: podocin-promoter driven diphtheria toxin receptor transgene; *n*=6 rats/group; *$P<0.05$, **$P<0.01$, ***$P<0.001$.

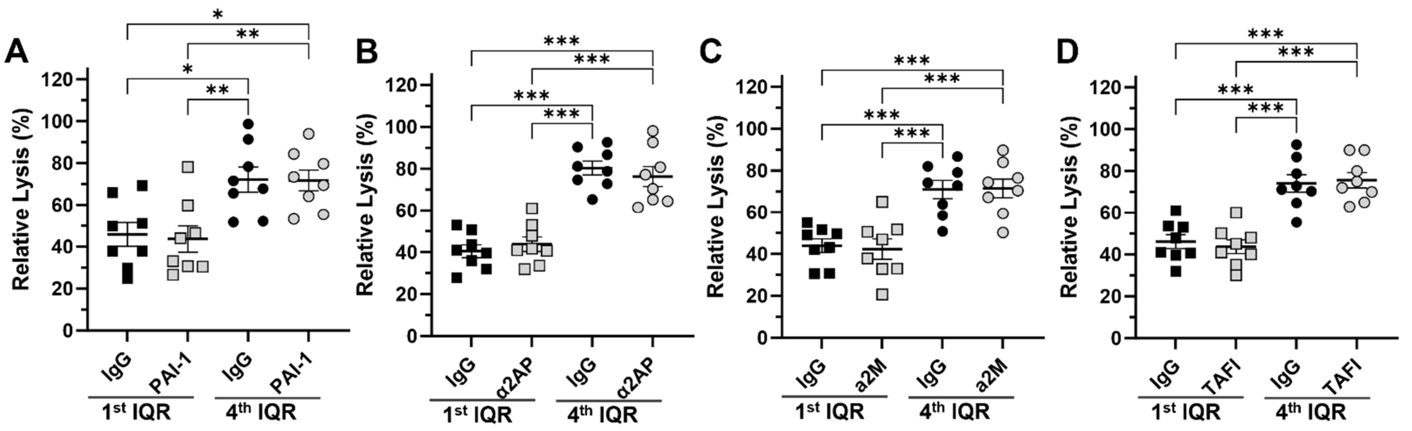


**Figure S6: Plasma Immunodepletion of Fibrinolytic System Proteins that were Increased Relative to Fibrin(ogen) in Clots formed from Hypoalbuminemic Plasma Failed to Improve Fibrinolysis.** (**A-D**) Hypoalbuminemia (1st IQR) significantly reduced fibrinolysis compared to normal albumin levels (4th IQR) in all conditions tested. Plasma immunodepletion of PAI-1 (**A**), α2AP (**B**), α2M (**C**), and TAFI (**D**) did not significantly alter fibrinolysis. 1st IQR: lowest albumin quartile *pDTR* rat plasmas; 4th IQR: highest quartile *pDTR* rat plasmas (normal); *pDTR*: podocin-promoter driven diphtheria toxin receptor transgene; IgG: non-specific isotype control antibody; PAI-1: plasminogen activator inhibitor-1; α2AP: α2-antiplasmin; α2M: α2-macroglobulin; TAFI: thrombin activatable fibrinolysis inhibitor; $n$=8 rats/group; *$P$<0.05, **$P$<0.01, ***$P$<0.001.

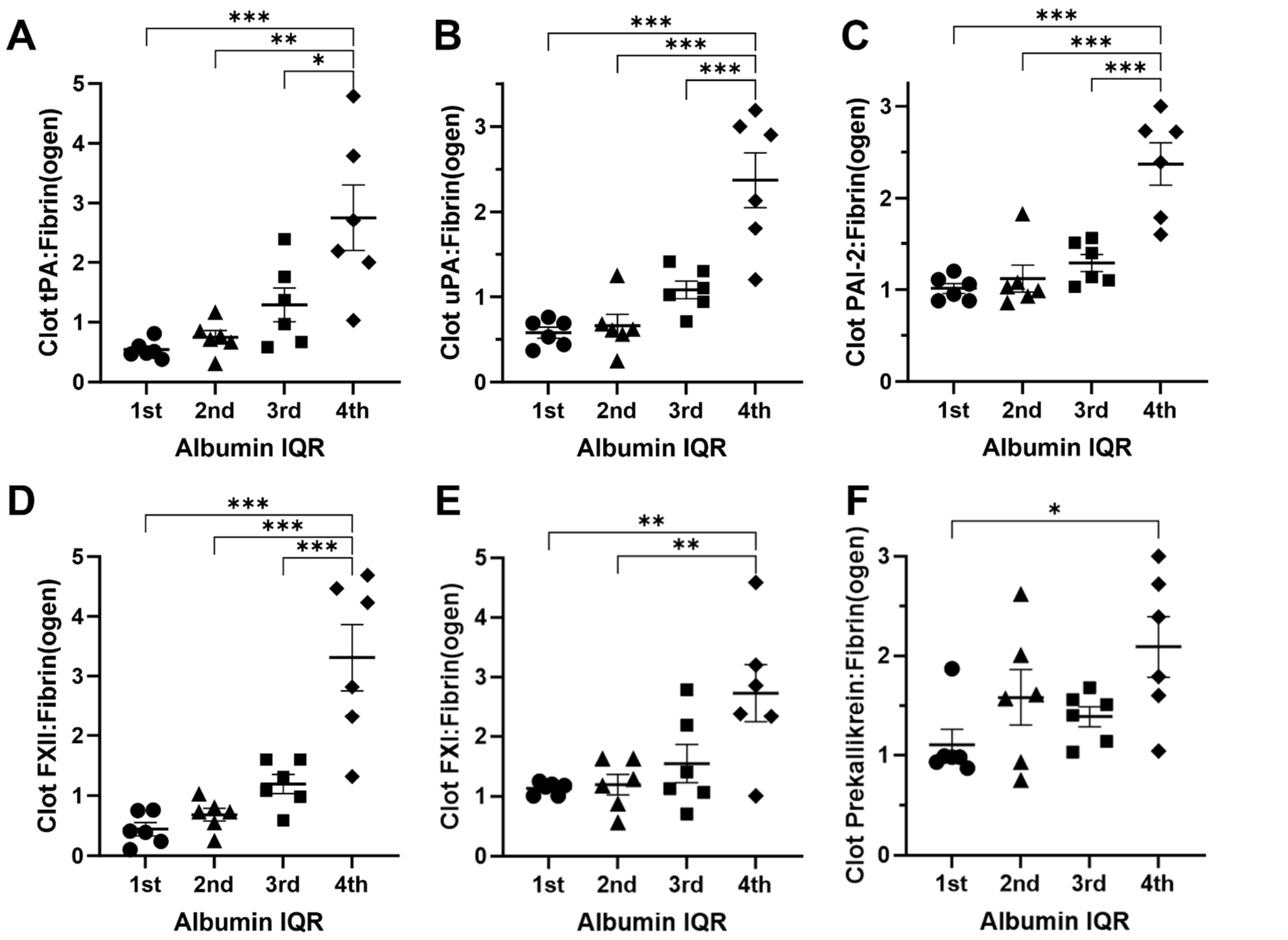


**Figure S7: Fibrinolytic System Proteins that Decreased Relative to Fibrin(ogen) in Clots formed from Hypoalbuminemic Plasma.** (**A-F**) Levels of tPA (**A**), uPA (**B**), PAI-2 (**C**), FXII (**D**), FXI (**E**), and Prekallikrein (**F**) were decreased relative to fibrin(ogen) in clots formed from hypoalbuminemic *pDTR* rat plasma. tPA: tissue-type plasminogen activator; uPA: urokinase; PAI-2: plasminogen activator inhibitor-2; FXII: factor XII; FXI: factor XI; IQR: interquartile range; *pDTR*: podocin-promoter driven diphtheria toxin receptor transgene; *n*=6 rats/group; **P*<0.05, ***P*<0.01, ****P*<0.001.

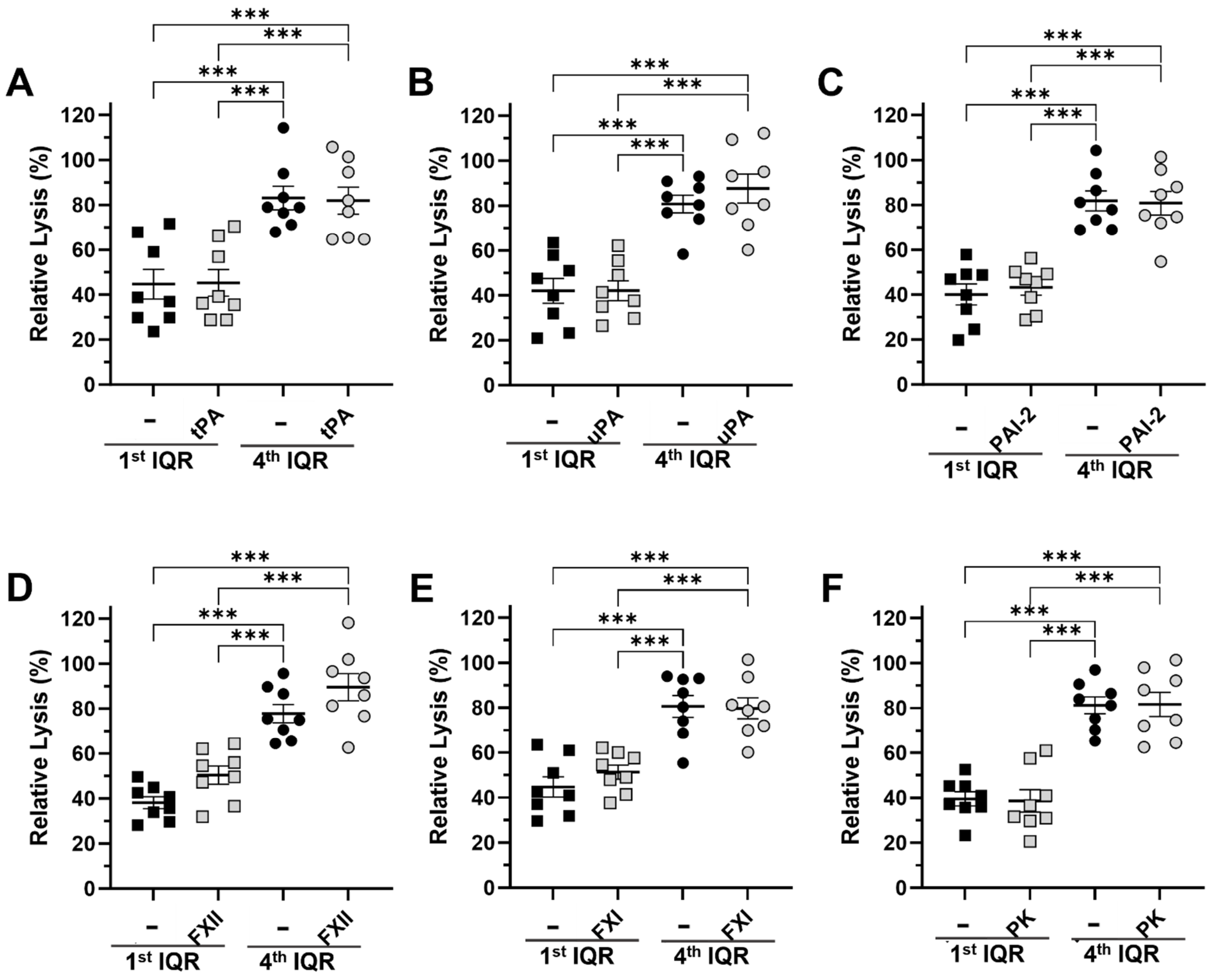


**Figure S8: Plasma Spiking with Fibrinolytic System Proteins that were Decreased Relative to Fibrin(ogen) in Clots formed from Hypoalbuminemic Plasma Failed to Improve Fibrinolysis.** (**A-D**) Hypoalbuminemia (1st IQR) significantly reduced fibrinolysis compared to normal albumin levels (4th IQR) in all conditions tested. Spiking tPA (**A**), uPA (**B**), PAI-2 (**C**), FXII (**D**), FXI (**E**), and Prekallikrein (**F**) into plasma did not significantly alter fibrinolysis. 1st IQR: lowest albumin quartile *pDTR* rat plasmas; 4th IQR: highest quartile *pDTR* rat plasmas (normal); *pDTR*: podocin-promoter driven diphtheria toxin receptor transgene; "–": buffer conrol; tPA: tissue-type plasminogen activator; uPA: urokinase; PAI-2: plasminogen activator inhibitor-2; FXII: factor XII; FXI: factor XI; PK: Prekallikrein; *n*=8 rats/group; ****P*<0.001.

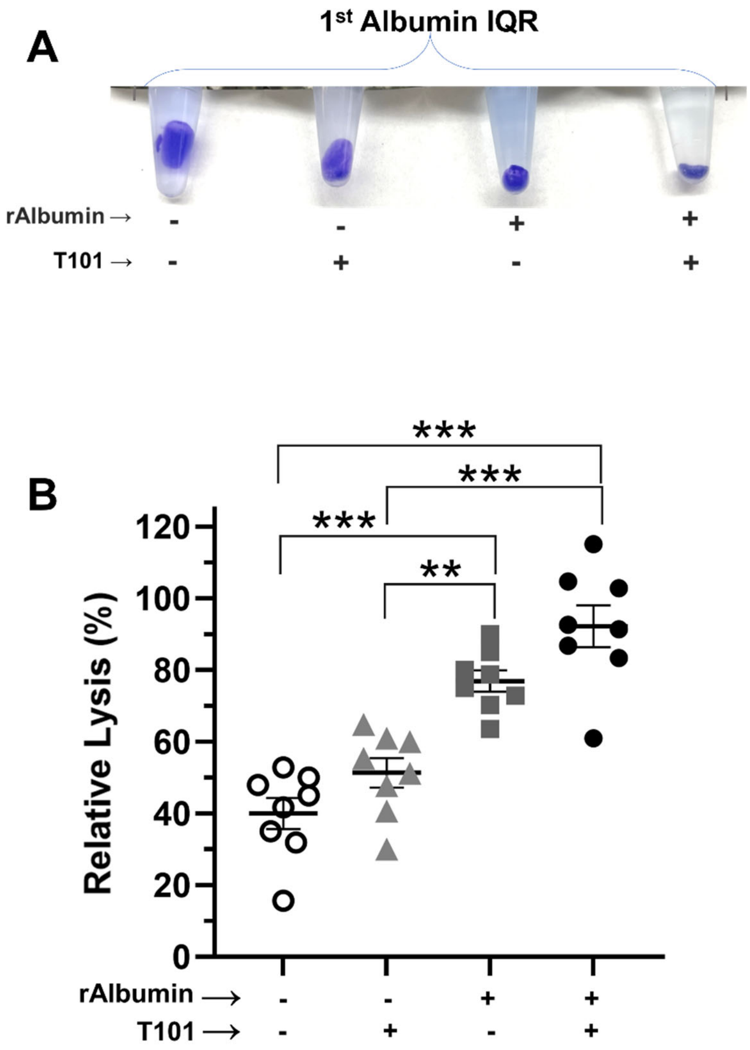


**Figure S9: Albumin-Dependent Fibrinolysis was Factor XIII Independent.** (**A**) Representative residual clots at the clot lysis endpoint (60 minutes) from 1st IQR hypoalbuminemic *pDTR*-nephrotic rat plasma without and with recombinant Albumin supplementation in the absence and presence of a factor XIII-specific inhibitor (T101; 20 μM). (**B**) Albumin supplementation significantly increased fibrinolysis whereas T101-mediated factor XIII inhibition did not significantly alter fibrinolysis. 1st IQR: lowest albumin quartile *pDTR* rat plasmas; rAlbumin: recombinant albumin; *n*=8 rats/group; ***P*<0.01, ****P*<0.001.

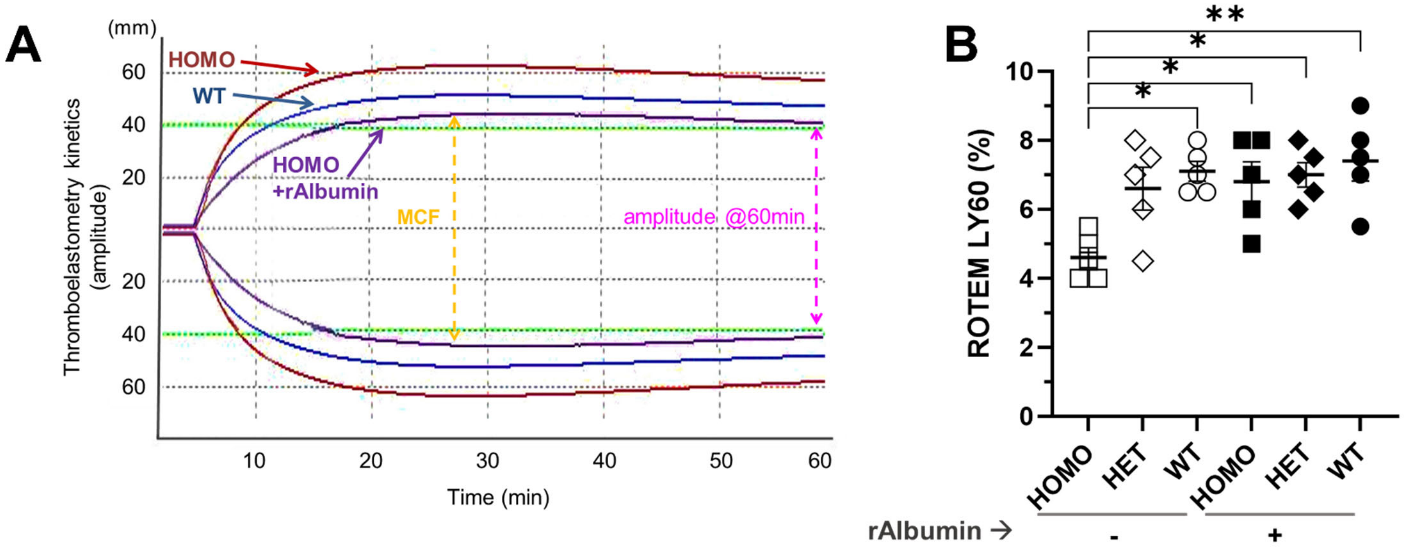


**Figure S10: Rotational Thromboelastometry Revealed a Hypofibrinolytic Defect in Albumin Knockout Rat Whole Blood.** (**A**) Representative ROTEM tracings from littermate wild-type and homozygous albumin knockout rats and homozygous blood supplemented with recombinant albumin. (**B**) %Lysis (LY60) determined by ROTEM demonstrated an albumin gene-dose effect and was corrected by recombinant albumin supplementation. WT: wild-type; HET: heterozygous; HOMO: Homozygous; rAlbumin: recombinant albumin; *n*=5 rats/group; **P*<0.05, ***P*<0.01.

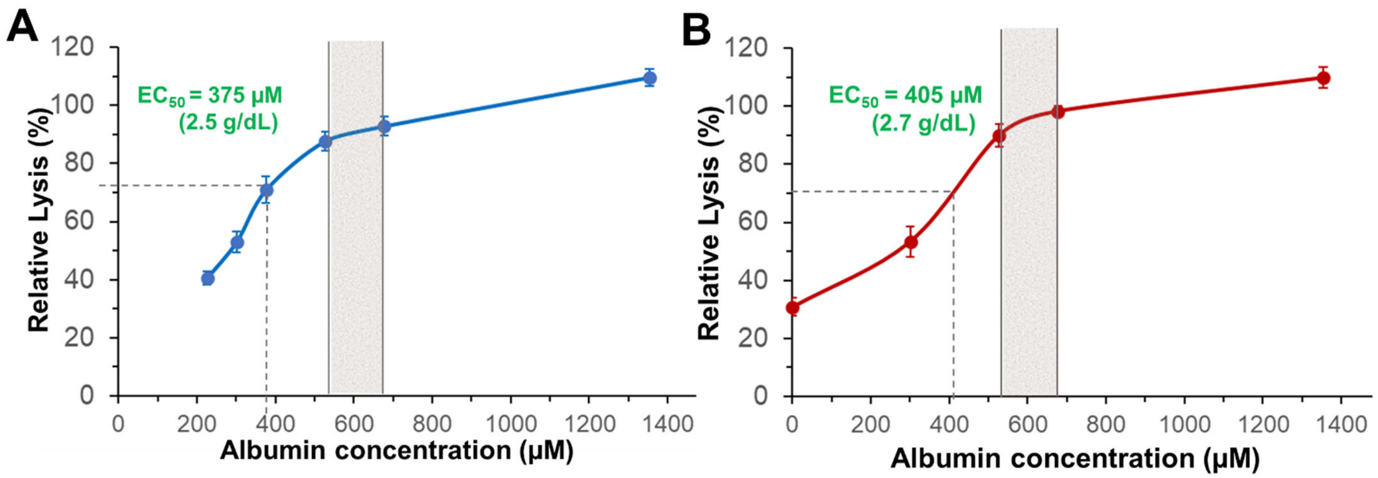


**Figure S11: Albumin Supports Fibrinolysis at Physiologic Concentrations.** Albumin dose-response curve in *pDTR* (**A**) and albumin knockout (**B**) rat plasma clots with half-maximal effective concentration ($EC_{50}$) estimates. Shaded area represents the healthy rat albumin reference range (3.5-4.5 g/dL; 526-677 µM). *n*=5-8/concentration point.
